# Single cell transcriptomics of human epidermis reveals basal stem cell transition states

**DOI:** 10.1101/784579

**Authors:** Shuxiong Wang, Michael L. Drummond, Christian F. Guerrero-Juarez, Eric Tarapore, Adam L. MacLean, Adam R. Stabell, Stephanie C. Wu, Guadalupe Gutierrez, Bao T. That, Claudia A. Benavente, Qing Nie, Scott X. Atwood

## Abstract

How stem cells give rise to human interfollicular epidermis is unclear despite the crucial role the epidermis plays in barrier and appendage formation. Here we use single cell-RNA sequencing to interrogate basal stem cell heterogeneity of human interfollicular epidermis and find at least four spatially distinct stem cell populations that decorate the top and bottom of rete ridge architecture and hold transitional positions between the basal and suprabasal epidermal layers. Cell-cell communication modeling through co-variance of cognate ligand-receptor pairs indicate that the basal cell populations distinctly serve as critical signaling hubs that maintain epidermal communication. Combining pseudotime, RNA velocity, and cellular entropy analyses point to a hierarchical differentiation lineage supporting multi-stem cell interfollicular epidermal homeostasis models and suggest the “transitional” basal stem cells are stable states essential for proper stratification. Finally, alterations in differentially expressed “transitional” basal stem cell genes result in severe thinning of human skin equivalents, validating their essential role in epidermal homeostasis and reinforcing the critical nature of basal stem cell heterogeneity.

## INTRODUCTION

Defining stem cell (SC) heterogeneity and its functional consequences in tissue homeostasis remains an open question in biology. In the skin, SCs reside in the basal compartment of the interfollicular epidermis (IFE) and in discrete compartments within ectodermal skin appendages – namely the pilosebaceous unit and sweat gland (Lu et al., 2012). Decades of work primarily from mouse studies have demonstrated the presence of multiple SC pools residing in various compartments of the pilosebaceous unit, including the bulge (Blanpain et al., 2004; Cotsarelis et al., 1990; Liu et al., 2003; Lyle et al., 1998; Morris et al., 2004; Trempus et al., 2003; Tumbar et al., 2004), hair germ (Greco et al., 2009; Ito and Kizawa, 2001; Ito et al., 2002; Rompolas et al., 2012), isthmus (Jensen et al., 2008; Nijhof et al., 2006; Snippert et al., 2010), junctional zone (Page et al., 2013), upper portion of the infundibulum (Jensen et al., 2008), and sebaceous gland (Cottle et al., 2013; Horsley et al., 2006; Jensen et al., 2009). In contrast to the pilosebaceous unit, basal SCs in the IFE are considered more homogenous and are thought to have one or two distinct subpopulations depending on the body site in mouse or human. However, a large degree of plasticity exists within the skin stem cell populations. Bulge or IFE stem cells can both form the pilosebaceous unit and sweat glands in response to inductive signals from the underlying dermis (Jones et al., 2007), suggesting that IFE stem cells may be more heterogenous than previously thought. Whether epidermal stem cells exist on a continuum and can equally respond to inductive signals or whether they occupy more stable states is unclear.

IFE self-renewal is thought to be achieved under homeostatic conditions by proliferation of SCs in the basal compartment, followed by transit amplification and terminal differentiation of the SC progeny (Watt, 1998). Early pedigree studies in mouse dorsal skin suggest that a single basal SC gives rise to transit amplifying (TA) cells with limited proliferation capacity that are destined to undergo terminal differentiation, coined the epidermal proliferative unit (EPU) (Mackenzie, 1975, 1997; Potten, 1974). Other models suggest a single population of committed progenitor cells that directly self-renew or differentiate (Clayton et al., 2007; Rompolas et al., 2016), a slower cycling stem cell population that gives rise to committed progenitor cells that directly differentiate (Mascre et al., 2012), or two independent stem cell populations that regenerate at different rates (Sada et al., 2016).

How human IFE self-renewal is achieved is unclear, primarily because of its complex architecture and absence of genetic and imaging tools. Using long-term fate mapping strategies enabled by a lentivirus-mediated gene transfer approach (Ghazizadeh et al., 2004), basal SCs appear to be dispersed along the basal compartment in human foreskin epidermis and that EPUs are present and capable of engaging in epidermal self-renewal in human xenografts (Ghazizadeh and Taichman, 2005). This gene transfer approach also enabled the observation that basal SCs do not preferentially occupy specific regions along the basal layer, as it was previously proposed (Lavker and Sun, 1982, 2000), but rather occupy locations throughout the basal compartment of human skin. However, whether each EPU arose from similar or distinct SC populations remains unclear as the widths and columns of the EPUs varied considerably depending on the originating site.

The advent of single cell RNA-sequencing (scRNA-seq) technologies has enabled the study of cellular heterogeneity, reconstruction of lineage hierarchies, inference of signaling networks, and has partially enabled the dissemination of functional SC roles in complex tissues. scRNA-seq has been successfully applied to normal human tissues including skin epidermis (Cheng et al., 2018) and dermis (Philippeos et al., 2018; Tabib et al., 2018); skin-related pathologies such as nasal polyps (Ordovas-Montanes et al., 2018), acne, alopecia, leprosy, psoriasis and granuloma annulare (Hughes et al., 2019); and various human cancers including melanoma (Tirosh et al., 2016), breast (Karaayvaz et al., 2018; Nguyen et al., 2018), and head and neck (Puram et al., 2017). In mice, scRNA-seq has revealed extensive functional heterogeneity in skin (Dong et al., 2018; Gupta et al., 2019; Mok et al., 2019; Salzer et al., 2018), hair follicles (Joost et al., 2016; Yang et al., 2017), and regenerative and non-regenerative wounds (Guerrero-Juarez et al., 2019; Joost et al., 2018). Despite these studies, epidermal stem cell heterogeneity of human IFE remains unresolved. To address this issue, we interrogated epidermal cell heterogeneity within human neonatal foreskin epidermis using droplet-enabled scRNA-seq and identify four spatially distinct basal stem cell subpopulations. Interrogation of the “transitional” basal subpopulations that spatially occupy both the basal and suprabasal layers indicate their essential role in epidermal homeostasis. Our findings argue against a single population of progenitor cells and suggest a more complex model of multiple epidermal stem cell transitions that maintain epidermal homeostasis.

## RESULTS

### Single cell transcriptome analysis reveals robust cellular heterogeneity in human neonatal epidermis

To define the cellular heterogeneity of human IFE, we isolated viable, single cells from discarded and de-identified human neonatal foreskin epidermis and subjected them to droplet-enabled scRNA-seq to resolve their individual transcriptomes (Figure 1A; Supplementary Figure 1; Supplementary Table 1; n = 5). We chose foreskin epidermis because it is composed of mostly IFE and contains few rudimentary skin appendages, such as hair follicles and sweat glands (Tuncali et al., 2005). We processed a total of 17,553 cells and performed quality control analysis on individual libraries using Seurat (Supplementary Figure 2) (Macosko et al., 2015). We used *Similarity matrix-based OPtimization for Single Cell* (SoptSC) to bioinformatically parse and analyze our data (Wang et al., 2019). The SoptSC algorithm is based on a cell-cell similarity matrix that can coherently perform many inference tasks – including unsupervised clustering, pseudotemporal ordering, cell lineage inference, cell-cell communication, and network inference. We used library three as a representative of our data sets for analysis, as it contained the highest median gene number per cell (3,104 median genes per cell). Library three contained 4,728 total cells where 4,598 met quality control standards and were further used for downstream bioinformatics analyses (Supplementary Figure 2, *Methods*). SoptSC clustered cells into eight distinct cell communities, corresponding to four distinct cellular cohorts, using two-dimensional elastic embedding (EE) (Figure 1B) (Carreira-Perpiñan, 2010). A cellular cohort is defined as a group of cell communities expressing similar known marker genes. Three-dimensional EE further demonstrated non-overlapping cell communities inferred by SoptSC (Supplemental Figure 3). By comparing gene expression profiles across cell communities, we characterized their putative cellular identities and proportions (Figure 1B). Basal SC communities BAS-I – BAS-IV represented approximately ∼2%, ∼5%, ∼6% and ∼17% of the entire population pool, respectively, and were enriched for known basal keratinocyte marker genes including *KRT14*, *KRT5*, *DSG3*, and *CDH3* (Figure 1C, D). Although known basal marker genes were able to distinguish between cellular identities (i.e. basal *vs*. granular keratinocytes), they were not sufficient to distinguish between basal clusters despite being clustered distinctly by SoptSC. Spinous communities SPN-I – SPN-II, representing approximately ∼30% and ∼21% of the entire population pool, respectively, showed heightened expression of *KRT1*, *KRT10*, *DSG1*, and *CDH1* that continued to be expressed in granular keratinocytes (Figure 1C, D). Similarly, spinous marker genes alone were not sufficient to distinguish between spinous clusters, even when they were clustered distinctly by SoptSC. Differentiated granular keratinocytes (GRN, ∼12% of total cells) expressed the differentiation gene markers *DSC1*, *KRT2*, *IVL*, *LOR*, and *TGM3*. SoptSC clustered melanocytes and Langerhans cells together into one single cluster (MEL/LAN, ∼7%). This cluster was enriched for the melanocytic markers *MITF* and *MLANA*, as well as the Langerhans markers *CD207* and *CD86*.

**Figure 1.**
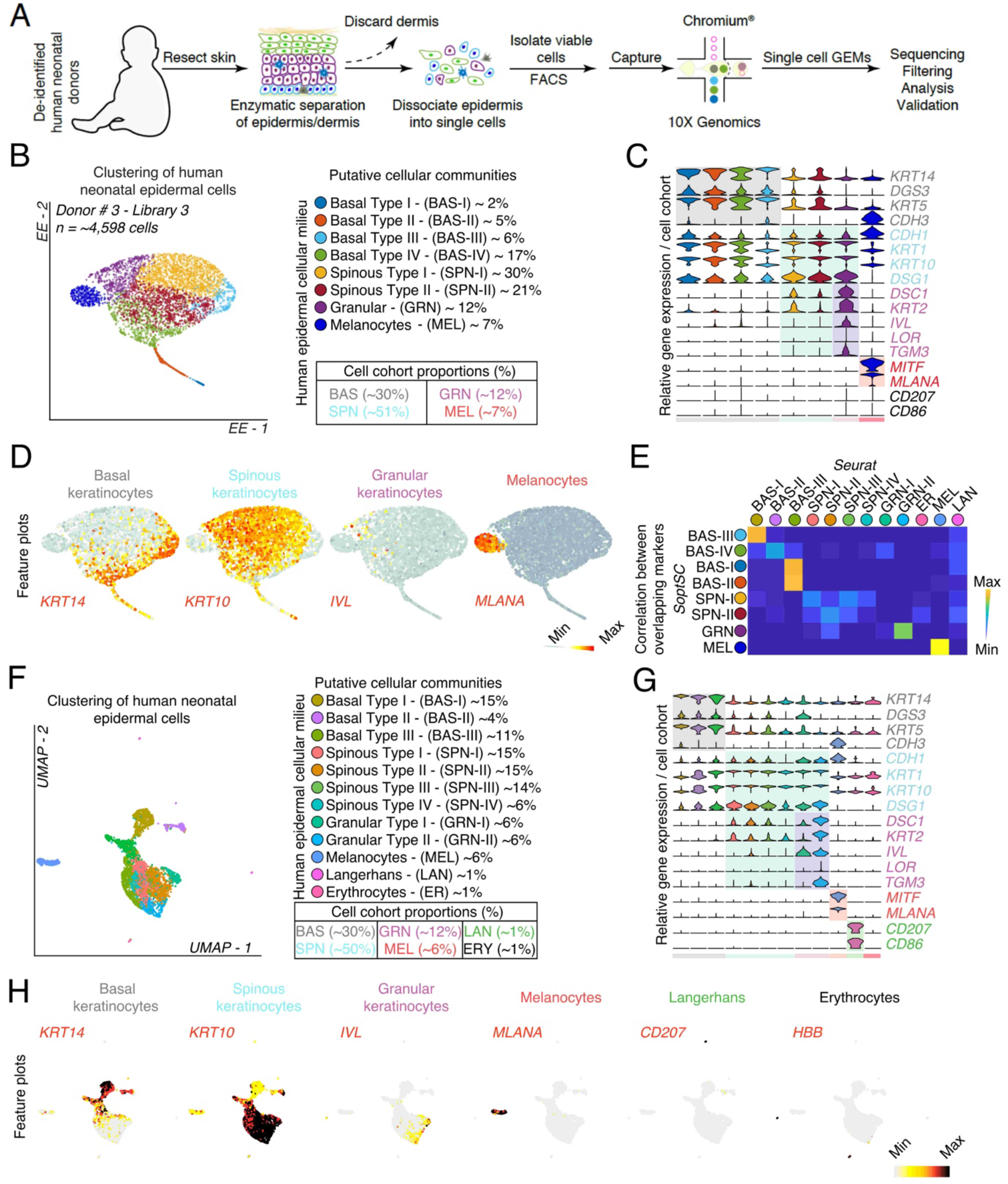
Defining human neonatal epidermal cell populations using single cell RNA-sequencing. **A)** Schematic of epidermal cell isolation from human neonatal foreskin. **B)** Clustering of 4,598 single cells isolated from Library 3 using SoptSC and displayed using Elastic Embedding (EE). Cell proportions from putative cellular communities are quantified on the right. **C)** Violin plots of relative gene expression of known epidermal marker genes split by cell cohorts. **D)** Feature plots showing expression of keratinocyte markers from basal, spinous, and granular layers, including melanocytes. **E)** Correlation between overlapping differentially expressed genes from SoptSC and Seurat. **F)** Clustering of 4,598 single cells isolated from Library 3 using Seurat and dimensionality reduction using UMAP. Cell proportions from putative cellular communities are quantified on the right. **G)** Violin plots of relative gene expression of known epidermal marker genes split by cell cohorts. **H)** Feature plots showing expression of keratinocyte markers from basal, spinous, and granular layers, including melanocytes, Langerhans cells, and erythrocytes.

To further corroborate the robustness of SoptSC, we compared its clustering performance with Seurat. To do this, we used the supervised clustering feature in Seurat, which identified similar cell communities as SoptSC (Figure 1F). For example, basal SC communities were grouped into three distinct communities – collectively representing ∼30% of the entire cell pool, whereas spinous and granular cellular cohorts represented ∼50% and ∼12% of the cell pool, respectively. Furthermore, Seurat clustered melanocytes and Langerhans cells separately based on expression of *MITF* and *MLANA* (MEL ∼6%) and *CD207* and *CD86* (LAN ∼1%), and identified a community composed of erythrocytes (ER ∼1%) based on expression of *HBB-2A* (Figure 1F). We also compared gene expression profiles across cell communities (Figure 1G). Dimensionality reduction strategies employed by Seurat and SoptSC remain generally congruent on clustering performance, cell type, and distribution of cell communities (i.e. number of overlapped markers between clusters) across all our individually sequenced libraries (Figure 1E; Supplementary Figure 4). In sum, our comparative analysis helps to reconcile the ability of SoptSC to unbiasedly cluster main cohorts and individual communities found in human neonatal epidermis and corroborates previous histological assessment of cell type and approximate proportion *in vivo* (Tuncali et al., 2005).

### Identification and spatial characterization of keratinocytes in human neonatal epidermis

To define novel genes associated with basal keratinocytes and human epidermis as a whole, we analyzed differentially expressed genes that define each cluster and found marker genes that provide more specific resolution for each cluster than widely used epidermal marker genes (Figure 2A-C; Supplementary Table 1). KRT14 immunofluorescence uniformly spans several layers in human neonatal epidermis with KRT10 staining beginning in the second layer and enriching in subsequent layers (Figure 2D). Basal cluster BAS-III is defined by expression of *ASS1*, *COL17A1*, and *POSTN*, where ASS1 and COL17A1 immunofluorescence staining of neonatal human epidermis shows enrichment between rete ridges, suggesting a specific zone of basal stem cells surrounding the papillary dermis (Figure 2E). The BAS-IV basal cluster is defined by expression of the gap junction gene *GJB2*, *KRT6A*, and *RHCG.* Immunofluorescence staining of GJB2 shows enrichment at the bottom of the rete ridges with some expression in the upper strata, whereas BAS-III cluster gene KRT19 shows enrichment at the bottom/side of the rete ridges (Figure 2F), reinforcing a specialized zone of basal stem cells that can be regenerated after partial-thickness wounding (Lin et al. 2019). The BAS-I and BAS-II basal clusters are enriched for cell cycle marker genes but are maintained even after cell cycle regression (Supplementary Figure 5). The topology of the keratinocyte sub-clusters in EE was maintained and cell community gene expression profiles remained congruent with one another, suggesting that cell cycle genes do not profoundly influence keratinocyte sub-clusters. BAS-I is defined by expression of *PTTG1*, *PLK1*, and *CDC20*, whereas BAS-II is defined by *RRM2*, *HELLS*, and *PCLAF* expression. Immunofluorescence staining of PTTG1, CDC20, PCLAF, and RRM2 show a transitional position within the epidermis where the cells occupy space between the basal and suprabasal layers (Figure 2G-H). Many of these cells are still adjacent to the basement membrane with the bulk of the cell body and nucleus residing either in the basal or suprabasal layers. These “transitional” basal cells appear to be in the process of delaminating from the basal layer, are spread heterogeneously across the epidermis, and may represent basal stem cells with a fluid cell fate.

**Figure 2.**
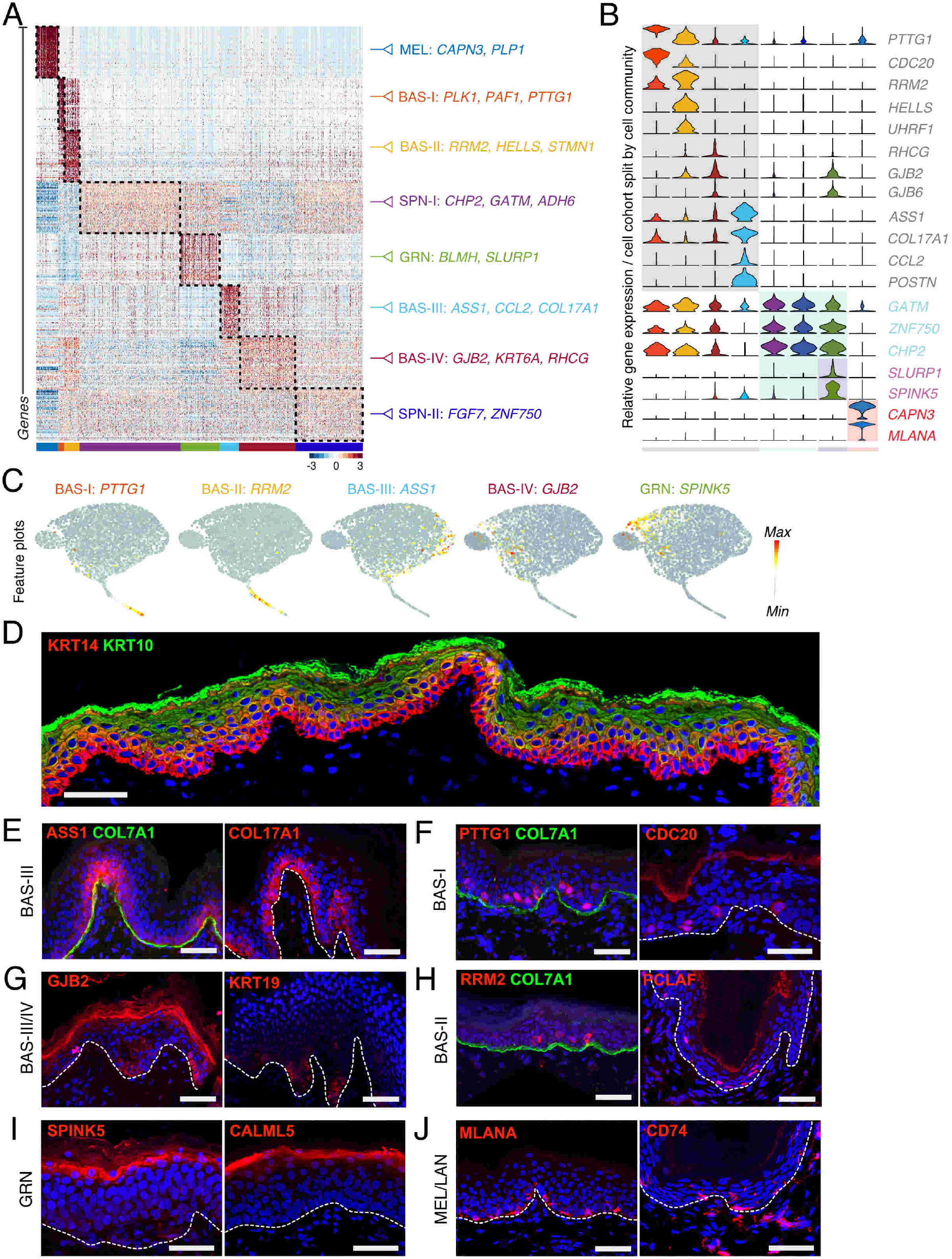
Differential gene expression and immunostaining of epidermal cells highlight basal stem cell heterogeneity. **A)** Heatmap showing top 300 differentially expressed genes per cluster. Dotted lines outline differentially expressed genes. Select marker genes are shown on the right. **B)** Violin plots of relative expression of marker genes split by cell cohorts and color-coded by cell community as in A. **C)** Feature plots showing expression of select basal and granular cell marker genes. **D)** Immunostaining of KRT14 (red), KRT10 (green), and DAPI (blue) in human neonatal skin. Scale bar 100µm. **E-J)** Immunostaining of differentially expressed proteins (red) from the **(E)** BAS-III cluster (ASS1 and COL17A1), **(F)** BAS-I cluster (PTTG1 and CDC20), **(G)** BAS-III/IV clusters (KRT19 and GJB2), **(H)** BAS-II cluster (RRM2 and PCLAF), **(I)** GRN cluster (SPINK5 and CALML5), and **(J)** the MEL (MLANA) and LAN (CD74) clusters in human neonatal skin. COL7A1 (green) and DAPI (blue) are co-stained. Dotted lines denote position of basement membrane where COL7A1 staining is absent. Scale bar 100µm.

We are also able to identify specific granular keratinocyte genes such as *SLURP1*, *SPINK5*, and *CALML5*, with the latter two showing robust protein enrichment in the granular layer of human neonatal epidermis (Figure 2I). Melanocyte (*MLANA*) and Langerhans (*CD74*) gene expression signatures also robustly mark both clusters, with protein expression highly restricted to these cell types (Figure 2J). However, we are not able to identify specific gene expression signatures that are strongly enriched only in the spinous keratinocytes or that spatially immunostain the spinous layer. The spinous keratinocytes are represented by two clusters – SPN-1 and SPN-2 – that both appear to segregate based on lack of basal- or granular-specific markers, suggesting they are at the beginning of a differentiation trajectory that ends with the granular fate and may not be a stable state by themselves.

### Cell-cell network inference reveals essential cross-talk between keratinocyte populations

A feature of single cell analysis is the ability to infer signaling networks within a cell and reconstruct potential cell-cell signaling interactions important in homeostasis, disease, and cancer (Kumar et al., 2018; Smillie et al., 2019). Having defined major cell communities in human neonatal epidermis, we sought to quantify potential cell-cell interactions (*P_i,j_*) using the probabilistic cell-cell network inference algorithm featured in SoptSC. Signaling relationships in SoptSC are calculated based on differential gene expression and co-variance of specific signaling pathway components. Signaling probabilities between cells are then defined and quantified based on the weighted expression of signaling pathway components between sender-receiver cell pairs inferred via expression of ligand-receptor pairs. We used a reference of known, literature-supported interactions from the JAK/STAT, NOTCH, TGF-β, and WNT signaling pathways and scored interactions between single cells (Supplementary Figure 6 and Supplementary Table 2) (Ramilowski et al., 2015; Subramanian et al., 2005).

The JAK/STAT pathway is comprised of four non-receptor tyrosine kinases that are typically activated by cytokine receptors and seven intracellular signaling substrates (Welsch et al., 2017). Abnormalities in this pathway can cause a number of skin-associated inflammatory disorders such as psoriasis, lupus erythematosus, atopic dermatitis, and alopecia areata. Our predicted JAK/STAT signaling interactions at both the single cell and community levels in epidermal keratinocytes show overwhelming activation (i.e. receiver cells) in granular cells (*P_i,j_φ* = 0.1) (Figure 3A). Although most of the aforementioned skin disorders are attributable to disruption of JAK/STAT signaling in immune cells, adult epidermis shows strong immunostaining for specific pathway components (JAK3, TyK2, and STAT2/3/4/6) in the *stratum granulosum* layer (Nishio et al., 2001). In addition, resident skin cells can produce cytokines that help promote skin barrier through cornification (IL-31), lipid envelope composition (IFN-g), and cell-cell adhesion (IL-1a) (Hanel et al., 2013), all processes that predominantly occur in the granular layer.

**Figure 3.**
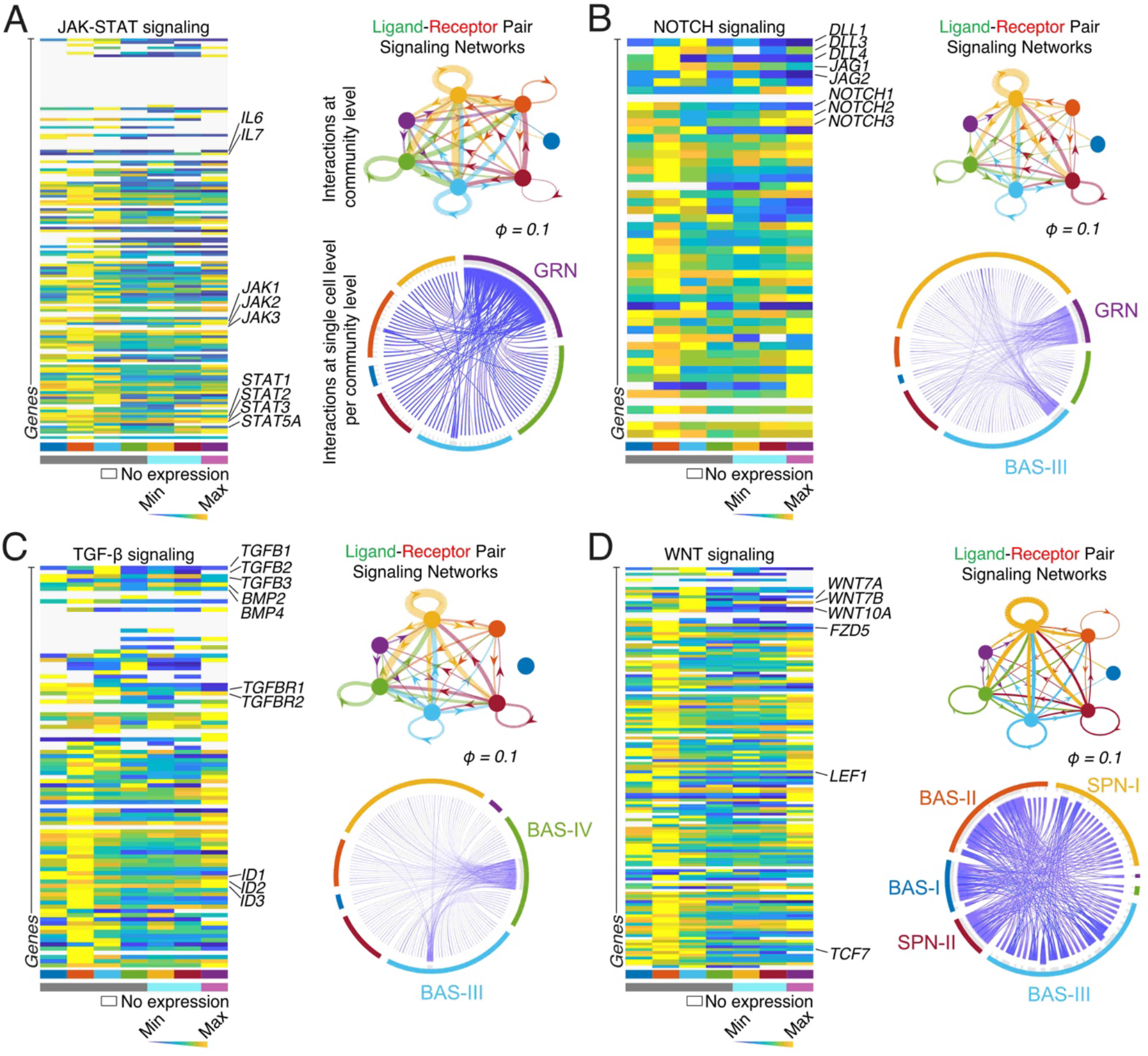
Cell-cell interaction modeling reveals auto- and paracrine signaling hubs essential for epidermal homeostasis. **A-D)** Cell-cell communication networks predicted for the **(A)** JAK-STAT, **(B)** NOTCH, **(C)** TGF-β, and **(D)** WNT signaling pathways. Left: Heatmap showing expression levels of cognate ligand-receptor pairs and their downstream targets for each respective pathway. Select genes are denoted. Colors correspond to cluster labels from Figure 4A and community labels from Figure 1B at bottom. Right (top): Cluster-to-cluster signaling interactions where edge weights represent sums of cell-cell interactions within and between clusters. Probability of potential cell-cell interactions (*P_i,j_ φ* = 0.1). Right (bottom): Cell-cell networks where edge weights represent the probability of signaling between cells.

NOTCH signaling comprises heterodimeric transmembrane receptors where cells expressing any of five NOTCH ligands bind and activate up to four adjacent NOTCH receptors on neighboring cells, thereby initiating cleavage of the intracellular domain and subsequent translocation into the nucleus to help facilitate target gene expression (Siebel and Lendahl, 2017). Psoriasis and all three major skin cancers are associated with disruption of NOTCH signaling. We observe activation of NOTCH signaling (*P_i,j_ φ* = 0.1) predominantly in the GRN and BAS-III clusters, with some activation in the SPN-II cluster (Figure 3B), suggesting these epithelial populations may be important receivers of NOTCH signaling and recapitulating known roles in cell fate specification, proliferation, and differentiation in these compartments (Rangarajan et al., 2001; Blanpain et al., 2006; Moriyama et al., 2008).

The TGF-β family consists of ligands from TGF-β, BMP, Activin, and GDF signaling pathways. These ligands bind to their respective kinase receptors to phosphorylate and activate downstream SMAD effectors to allow their translocation into the nucleus and help facilitate target gene expression (Heldin and Moustakas, 2016). TGF-β also serves as a tumor suppressor, where disrupted TGF-β fuels tumor heterogeneity and drug resistance in squamous cell carcinoma (Oshimori et al., 2015). We observe strong activation of this pathway (*P_i,j_ φ* = 0.1) in both the BAS-III and BAS-IV clusters (Figure 3C), recapitulating its role in suppressing basal cell proliferation through the TGF-β and activin ligands (Bamberger et al., 2005; Mou et al., 2016) and suggesting that these populations are uniquely responsive to the TGF-β family.

A host of secreted WNT ligands differentially bind to ten distinct Frizzled receptors during canonical WNT signaling to activate Dishevelled, subsequently leading to β-catenin stabilization and translocation into the nucleus to turn on WNT-target genes (Clevers, 2006; Clevers and Nusse, 2012; Nusse and Clevers, 2017). WNT signaling is a pleotropic signaling pathway heavily involved from the earliest stages of skin development where it helps specify the ectoderm down the skin epithelium lineage (Wilson et al., 2001) and helps specify skin appendages (Kratochwil et al., 1996). WNT signaling is also involved in adult skin homeostasis where overexpression or loss of WNT/beta-catenin results in a variety of intra- and interfollicular epidermis phenotypes (Andl et al., 2002; Choi et al., 2013; Lim and Nusse, 2013; Xu et al., 2017; Zhang et al., 2009). Our data suggests that WNT signaling appears to be active in most of the basal and spinous populations (*P_i,j_ φ* = 0.1), with the majority of receiver cells being present in BAS-I and BAS-II (Figure 3D). Basal SCs in IFE of glabrous skin form an autocrine mechanism important for SC self-renewal, requiring WNT/β-catenin signaling to proliferate and, at the same time, producing and secreting long-range WNT inhibitors to promote differentiation (Lim et al., 2013). Deletion of β-catenin in the IFE of adult mice leads to a significant decrease in proliferation, suggesting that the WNT/β-catenin signaling contributes to progenitor cell proliferation under homeostatic conditions (Choi et al., 2013). In sum, our cell-cell network inference suggests that epidermal cell communities in human neonatal epidermis communicate through autocrine and paracrine signaling hubs and can recapitulate many of the reported skin-dependent roles of these major signaling pathways.

### Cellular entropy and RNA velocity predict the likelihood of cell state transitions and pseudotemporal directionality

Existing tools can enable the reconstruction of differentiation trajectories and therefore be used as a proxy to study developmental processes (Qiu et al., 2017; Schiebinger et al., 2019; Trapnell et al., 2014). To infer pseudotemporal cellular trajectories, SoptSC orders cells from the original cell-cell graph by calculating a collection of distances between an initial cell and all other surrounding cells. It then performs cell identity ordering and condenses individual cells into their respective larger clusters to generate a pseudotemporal trajectory. Basal keratinocytes undergo terminal differentiation into granular keratinocytes expressing the structural protein Involucrin (Watt, 1983). This stratification program occurs *in vivo* during homeostasis and in response to injury, and can be recapitulated in 2D and 3D *in vitro* culture systems (Bikle et al., 2012; Oh et al., 2013). During differentiation, keratinocytes progressively lose expression of basal markers, while they concomitantly begin to express suprabasal markers (Eckert and Rorke, 1989). This biological observation enabled us to make a direct comparison and accurately assess the modeled pseudotime trajectory inferred by SoptSC. We regressed MEL/LAN communities and used the remaining epithelial cohorts composed of BAS, SPN, and GRN communities to model a pseudotemporal trajectory of basal keratinocyte differentiation (Figure 4A). SoptSC unbiasedly reconstructed a putative BAS-SPN-GRN keratinocyte differentiation trajectory (Figure 4B). As expected, SoptSC placed basal keratinocytes expressing *KRT14* at the beginning, with average expression levels of *KRT14* declined towards the trajectory terminus, whereas cells expressing the terminal differentiation gene *TGM3* distributed differentially along the trajectory, displaying heightened average expression levels towards the trajectory terminus (Figure 4C). These results suggest that the unbiased pseudotemporal ordering inferred by SoptSC is concordant with the known epidermal differentiation-stratification program in skin and represents a putative BAS-SPN-GRN differentiation trajectory in human neonatal epidermis.

**Figure 4.**
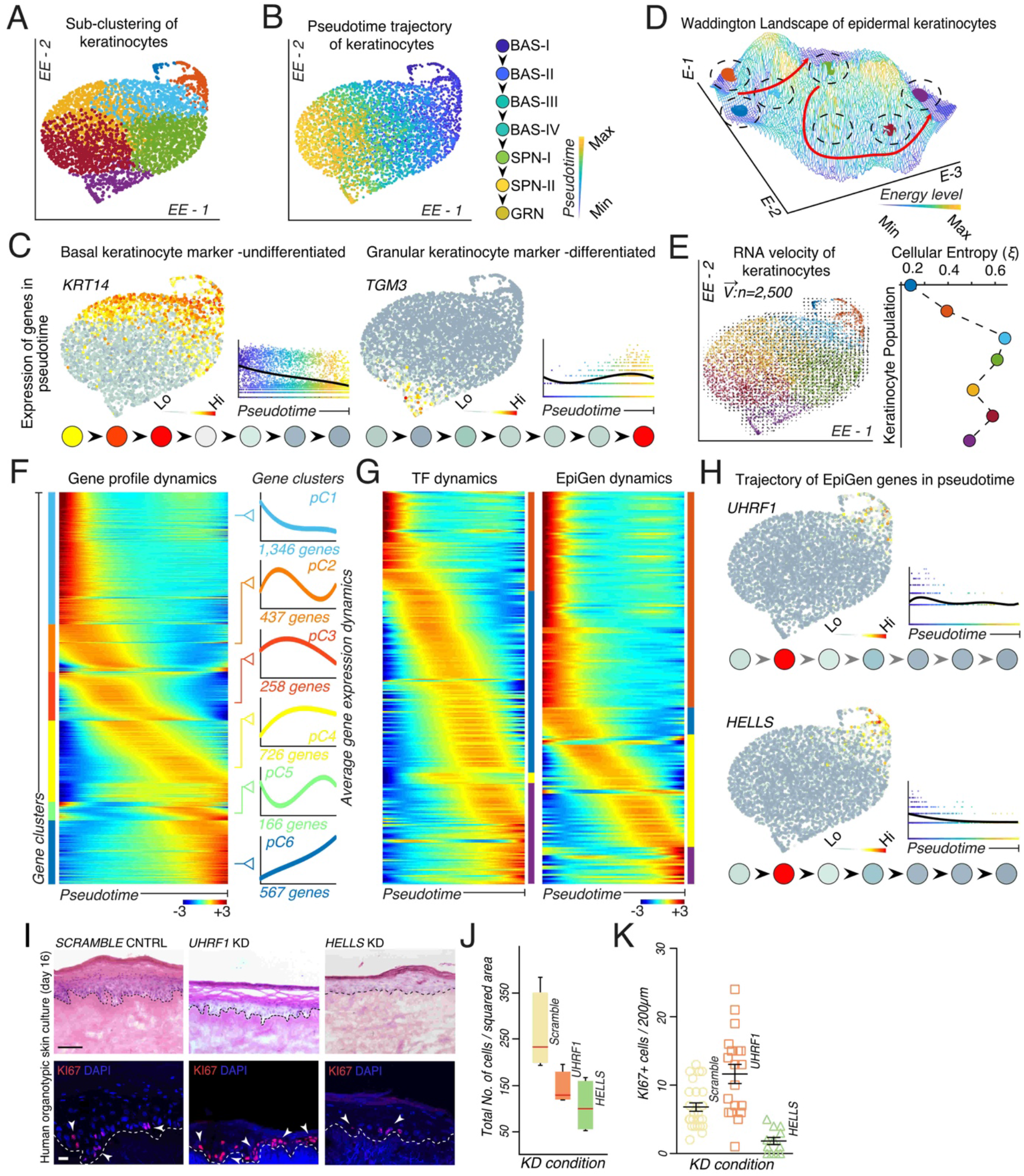
Pseudotime, lineage, and entropy analysis reveals keratinocyte differentiation trajectories and highlight the importance of epigenetic modifiers in epidermal homeostasis. **A)** Subclustering of epidermal keratinocytes using SoptSC and displayed using EE. **B)** Pseudotime inference of epidermal keratinocytes displayed using EE. Cell lineage inference displayed on the right. **C)** Feature plots showing expression of basal (KRT14) and granular (TGM3) genes and their expression along pseudotime. Expression along the epidermal cell lineage as determined in B displayed at the bottom. **D)** Cellular Entropy (*ξ*) of epidermal keratinocytes plotted in a Waddington landscape. Clusters color-coded as in A and displayed at their center position within the Waddington landscape. Red arrows show the predicted low energy path along differentiation. **E)** RNA velocity of epidermal keratinocytes overlaid on EE space using 2,500 vectors. Quantification of Cellular Entropy. Cells or clusters color-coded as in A. **F)** Rolling wave plot showing pseudotime-dependent gene expression dynamics. Average gene expression dynamics of each cluster pattern (pC1-6) shown on the right and independently color-coded. Number of genes in each cluster as labelled. **G)** Rolling wave plots showing pseudotime-dependent gene expression dynamics of transcription factors (TF) and epigenetic modifiers (EpiGen). Genes color-coded as in A on the right of each plot. **H)** Feature plots showing expression of *UHFR1* and *HELLS* and their expression along pseudotime. Expression along the epidermal cell lineage as determined in B displayed at the bottom. **I)** Knock-down (KD) of *UHFR1* and *HELLS*, with a scramble control, in primary human neonatal keratinocytes seeded on human devitalized dermis and grown at the air-liquid interface for 16 days. Hematoxylin and eosin (H&E) staining shown at top. Scale bar 100µm. KI67 (red) and DAPI staining shown at the bottom. Scale bar 50µm. Dotted lines denote the position of the basement membrane. Arrow heads pointing to some of the KI67+ cells. n = 6 each condition. **J)** Quantification of the total number (No.) of cells per squared area. K) Quantification of KI67+ cells per 200µm area along the basement membrane.

Next, we asked if the epithelial communities exist on a continuum or have distinct cellular states. Previous studies have suggested that *in silico* differentiation potency and plasticity of single cells can be approximated by computing the signaling promiscuity of a cell’s transcriptome (Teschendorff and Enver, 2017). We developed a similar algorithm to calculate the Cellular Entropy (*ξ*) of single cells called *Cellular Entropy Estimator (CEE)* – now a feature that has been incorporated into SoptSC (see Methods). CEE allows us to estimate the likelihood a single cell will transition from one cellular state to another. We applied CEE to estimate the Cellular Entropy in BAS, SPN, and GRN communities and represented their transition likelihoods in a Waddington landscape, where a “valley” represents a region of low Cellular Entropy (i.e. low likelihood of transition into a new state) and a “mountain” represents a region of high Cellular Entropy (i.e. high likelihood of transition into a new state) (Figure 4D). Epidermal communities in aggregate displayed distinct entropy values, with BAS-III and BAS-IV having the highest probability of transitioning to a new state (*ξ_BAS-III_ = ∼0.65; ξ_BAS-IV_ = ∼0.60*). As RNA velocity can estimate the future state of cells by analyzing spliced and unspliced variants of mRNA in single cell data (La Manno et al., 2018), we reasoned that combining Cellular Entropy with RNA velocity would predict the transition potency and directionality of a cell or group of cells (i.e. the likelihood of transitioning and its transition directionality). Indeed, we observed that BAS-III and BAS-IV displayed larger velocity vectors pointing toward a spinous cluster (SPN-I), as expected (Figure 4E). SPN-I, having a high entropy value (*ξ_SPN-II_ = ∼0.58*), also displayed refined velocity vectors toward GRN. *RRM2-* and *PTTG1*-positive cells had low Cellular Entropy values (*ξ_BAS-I_ = ∼0.20;ξ_BAS-II_ = ∼0.40*) and lacked refined velocity vectors, suggesting that these cells may represent a steady state.

### The BAS-II epigenetic modifiers HELLS and UHRF1 affect basal keratinocyte proliferation

To determine what genes change along our modeled pseudotime trajectory, we identified pseudotime-dependent gene expression changes and discovered a total of ∼3,500 differentially expressed genes (DEGs) along the putative BAS-SPN-GRN differentiation trajectory (Figure 4F and Supplementary Table 3). These DEGs segregated differentially into six distinct clusters (pC1-pC6) according to their average gene expression dynamics along pseudotime. We focused our attention on transcription factors (TFs) and epigenetic modifiers (EpiGens), given their roles in controlling cell states during differentiation of keratinocytes (Botchkarev et al., 2012). Using this approach, we identified TFs previously implicated in epidermal homeostasis and differentiation, including *MAFB* (Lopez-Pajares et al., 2015; Miyai et al., 2016), *TP63* (Mills et al., 1999)*, CEBPA/*B (Lopez et al., 2009)*, KLF4* (Segre et al., 1999), *GRHL3* (Ting et al., 2005), *GATA3* (de Guzman Strong et al., 2006), and *OVOL1* (Lee et al., 2014) (Figure 4G, Supplementary Figure 7, and Supplementary Table 3). EpiGens included factors involved in DNA methylation such as *DNMT1* and its co-factor *UHRF1* (Sen et al., 2010), and covalent histone modifications such as *KDM6B* (Sen et al., 2008) and *HDAC1/2* (LeBoeuf et al., 2010). We also identified Polycomb component members, including *EZH1/2* (Ezhkova et al., 2009), *JMJ* (Mejetta et al., 2011), and *CBX4* (Mardaryev et al., 2016); the ATP-dependent chromatin remodelers *SMARCA4* (Indra et al., 2005) and *CDH4* (Kashiwagi et al., 2007); and the higher-order chromatin remodeler *SATB1* (Fessing et al., 2011) (Figure 4G, Supplementary Figure 8, and Supplementary Table 3).

Our analysis identified *HELLS* with heightened expression at the beginning and decreased expression along the pseudotime trajectory. HELLS is a SNF2-like helicase, known for its role in silencing chromatin regions via interaction with DNA methyltransferases (Ren et al., 2019) and functions during development and senescence (Geiman et al., 2001; Keyes et al., 2011). *HELLS*, along with *UHRF1*, are target genes downstream of the Retinoblastoma (RB) pathway, which is mediated by E2F transcription factors, and have been implicated in recruitment of DMNT3 and DMNT1, respectively (Benavente et al., 2014; Jung et al., 2017). Although their role in epidermal homeostasis in unclear, *HELLS* and *UHRF1* are both differentially expressed in the transitional basal BAS-II community (Figure 4H). To assess the role of *HELLS* in epidermal homeostasis, we knocked down (KD) *HELLS* in primary human neonatal keratinocytes using viral transduction and then seeded the genetically modified primary keratinocytes on top of devitalized human dermis to generate human skin equivalents (Supplementary Figure 9). Histopathological assessment of *HELLS* KD organotypic cultures show decreased epidermal thickness, a significant reduction in total numbers of cells per squared area, and a significant decrease in KI67-positive cycling cells compared to *SCRAMBLE* control (Figure 4I-K). *UHRF1* KD organotypic cultures also show decreased epidermal thickness, reduced total number of cells, and concomitant increase in KI67-positive cells (Figure 4I-K). The *UHRF1* KD phenotype is similar to *DMNT1* KD organotypic skin cultures, where epidermal thickness are reduced and an increase in G2/M phase cells are seen at the expense of S phase (Sen et al., 2010). Taken together, our functional experiments reveal the importance of the epigenetic modifiers *HELLS* and *UHRF1* are important for the transitional basal cells to regulate human epidermal homeostasis.

### Basal stem cell heterogeneity suggests hierarchically ordered stem cell pools contribute to human epidermal differentiation

Previous studies have suggested functional heterogeneity within the IFE that has led to four distinct models of how this compartment is formed: 1) a single committed progenitor population that directly self-renews or differentiates (Clayton et al., 2007; Rompolas et al., 2016), one stem cell population that gives rise to 2) TA cells or 3) committed progenitors that directly differentiate (Potten, 1974; Mascre et al., 2012), or 4) two stem cell populations that regenerate at different rates (Sada et al., 2016). To assess functional heterogeneity in the basal compartment and address which model our data supports, we sub-clustered both the KRT14-high expressing keratinocytes regardless of their origin and the four main basal populations using both SoptSC and Seurat (Figure 5A-C, Supplementary Figure 10, and Supplementary Table 4). Sub-clustering the four main basal populations provided better resolution of cells expressing differentiation markers and thus likely transitioning into the differentiation state, whereas sub-clustering KRT14-high expressing cells resulted in sub-sampling the existing basal subpopulations with an absence of differentiating cells. After sub-clustering, 7 subpopulations emerged that largely split the BAS-III and BAS-IV clusters (Figure 5C). The transitional basal clusters (BAS-I and BAS-II) remained largely intact, further supporting their low Cellular Entropy values and robust steady state (Figure 4E). Pseudotime and cell lineage inference analysis indicates a largely linear differentiation trajectory that is supported by RNA velocity (Figure 5D-E). The bifurcation in the cell lineage inference seems to be the result of cellular trajectories going from kBAS-III.3 to kBAS-III.1 subpopulations as seen in the RNA velocity plot (Figure 5E). Tracking *KRT14*/*KRT10* gene expression in the kBAS subpopulations as a proxy for differentiation status shows a gradual decrease in *KRT14* gene expression over pseudotime with little to no change in *KRT10* expression, further supporting a linear temporal trajectory towards differentiation that occurs before commitment to differentiation (Figure 5F). When pairing the *KRT14*/*KRT10* expression with a differentiation score that tracks expression of GRN-specific genes, we observe commitment to differentiation occurring in the kBAS-IV.2 subpopulation at the end of the cell lineage inference and pseudotime (Figure 5G). Incidentally, the kBAS-IV.2 cluster most resembles the Ivl-CreER+ committed progenitors in murine epidermis (Mascre et al., 2012), with their Krt14-CreER+ stem cells most resembling the kBAS-I cluster (Figure 5H and Suppmentary Figure 11). In addition, non-label retaining stem cells with high proliferative capacity in murine epidermis from Krt5-tTa; pTRE-H2B-GFP; Krt14-CreER; Rosa-tdTomato mice (Sada et al., 2016) overlap with kBAS-I and Krt14-CreER+ stem cell datasets, whereas the label retaining stem cells with low proliferation capacity most resemble the KBAS-III.1 subpopulation that is also enriched for the basal stem cell marker *ITGB1* (Figure 5H and Supplementary Figure 11). These data, coupled with pseudotime and lineage inference indicating that the basal populations are hierarchically ordered (Figure 4B and 5D) and RNA velocity indicating BAS-III and BAS-IV can both contribute to spinous cells (Figure 4E), suggest that multiple stem cell populations contribute to differentiation and agree with models presented by the Blanpain and Tumbar groups that describe multiple stem cell pools with different proliferation capacities (Mascre et al., 2012; Sada et al., 2016).

**Figure 5.**
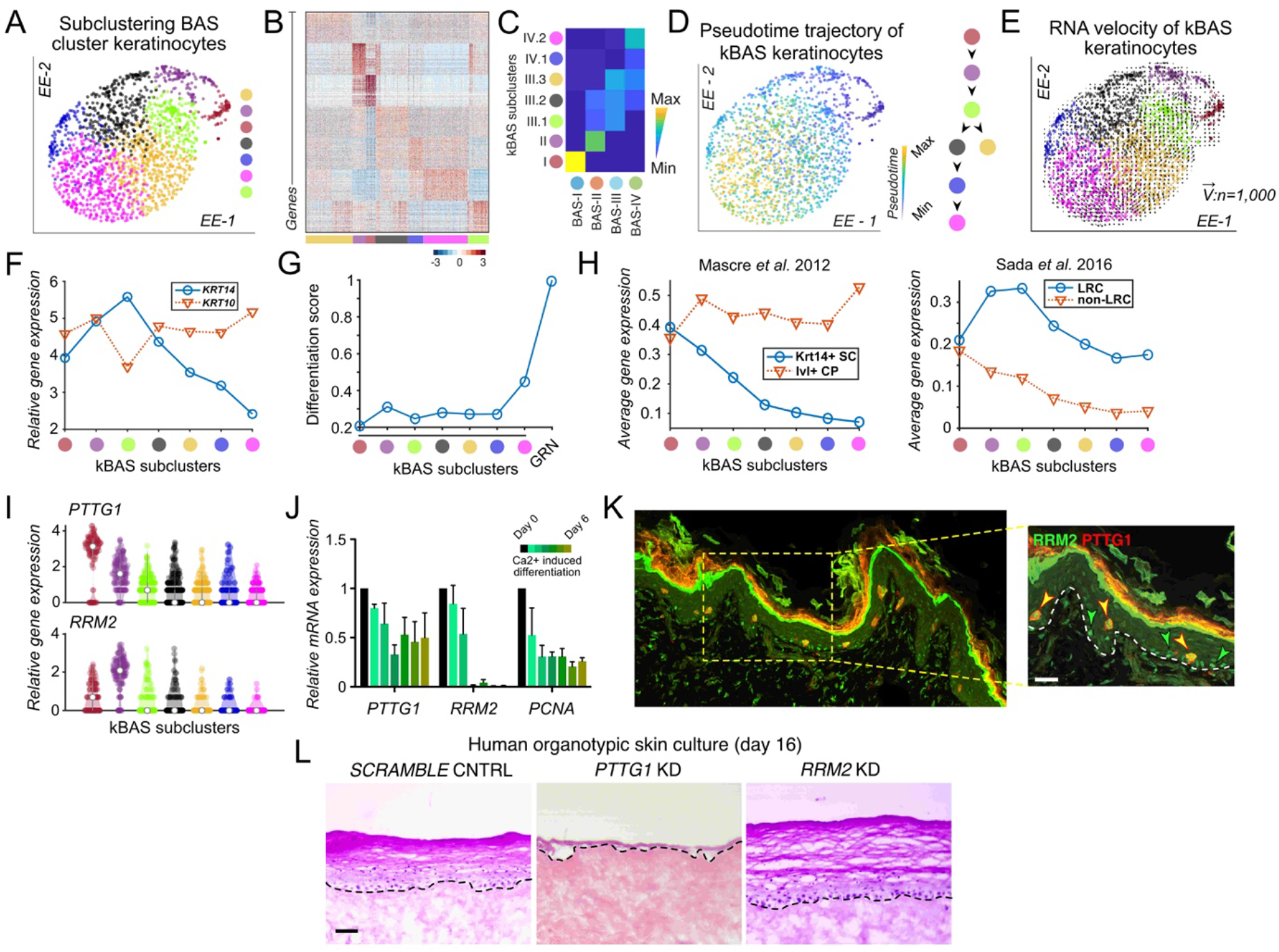
Subclustering basal stem cells clarify models of interfollicular epidermal differentiation and point to the BAS-I cluster as essential to epidermal homeostasis. **A)** Subclustering of basal cluster keratinocytes (BAS-I – BAS-IV) using SoptSC and displayed using EE. **B)** Heatmap showing top 100 differentially expressed genes per cluster. Cells are color-coded as in A at the bottom. **C)** Correlation between overlapping differentially expressed genes from basal clusters from Figure 3A and subclustering of basal clusters (kBAS) in A. Numbering of kBAS clusters corresponds to basal clusters in Figure 3A. **D)** Pseudotime inference of basal cluster keratinocytes displayed using EE. Cell lineage inference displayed on the right. **E)** RNA velocity of epidermal keratinocytes overlaid on EE space using 1,000 vectors. Cells are color-coded as in A. **F)** Relative gene expression of KRT14 and KRT10 across the kBAS subclusters along pseudotime. **G)** Differentiation score of each kBAS subcluster using average gene expression of differentially expressed GRN genes. **H)** Average gene expression of each kBAS subcluster using genes specific to (left) Krt14-CreER+ (Krt14+) stem cells (SC) or Ivl-CreER+ (Ivl+) committed progenitors (CP) from Mascre et al., 2012, or (right) label-retaining stem cells (LRC) or non-label-retaining stem cells (non-LRC) from Sada et al., 2016. **I)** Violin plot of relative gene expression of *PTTG1* and *RRM2* across the kBAS subclusters. **J)** Relative mRNA expression of *PTTG1*, *RRM2*, and *PCNA* along Ca^2+^-induced differentiation of primary human keratinocytes. n = 3. **K)** Co-immunostaining of PTTG1 (red) and RRM2 (green) in human neonatal skin. Green arrow heads highlighting RRM2-only expressing cells, whereas yellow arrow heads pointing to PTTG1-RRM2 double-positive cells. White dotted line denotes the position of the basement membrane. Scale bar 50µm. **L)** Knock-down (KD) of *PTTG1* and *RRM2*, with a scramble control, in primary human neonatal keratinocytes seeded on human devitalized dermis and grown at the air-liquid interface for 16 days. Hematoxylin and eosin (H&E) staining shown. Dotted lines denote the position of the basement membrane. Scale bar 100µm. n = 3 each condition.

We further profiled the kBAS subpopulations by assessing gene expression markers that encompass both transitional basal populations (BAS-I = kBAS-I and BAS-II = kBAS-II) and encompass the Krt14-CreER+ stem cells and non-label retaining cells with high proliferative capacity. kBAS-I can be defined primarily by expression of Pituitary Tumor Transforming Gene 1 (*PTTG1*), a proto-oncogene that is involved in controlling keratinocyte proliferation, early stages of differentiation, cell growth, and is overexpressed in psoriasis (Ishitsuka et al., 2013; Ishitsuka et al., 2012). kBAS-II can be defined by expression of Ribonucleotide Reductase 2 (*RRM2*), which controls the biogenesis of dNTPs and is overexpressed in skin cancer (Aird et al., 2013). Expression of *PTTG1*, *RRM2*, and fellow kBAS-II gene *PCNA* decreased in keratinocytes during Ca^2+^-induced differentiation (Figure 5J). In addition, co-immunofluorescence staining of the proteins showed RRM2-only expressing cells largely restricted to the basal layer, whereas double-positive RRM2 and PTTG1 cells occupying both basal and suprabasal positions, further reinforcing our observation that these genes are expressed in undifferentiated keratinocytes (Figure 5K). To determine the role of *PTTG1* in epidermal maintenance, we performed *PTTG1* KD in primary human neonatal keratinocytes using viral transduction and then seeded the genetically modified primary keratinocytes on top of devitalized human dermis to generate human skin equivalents (Supplementary Figure 9). *PTTG1* KD led to a severe epidermal phenotype, primarily characterized by disruption of the basal layer and differentiation program compared to scramble shRNA controls, indicating its essential role in basal stem cell maintenance and epidermal homeostasis (Figure 5L). On the other hand, *RRM2* KD did not appear to play a substantial functional role in epidermal homeostasis in human skin equivalents (Figure 5L), suggesting that the RRM2 promoter may be a promising candidate for exquisite Cre expression for the interrogation of the kBAS-II population.

## DISCUSSION

Functional heterogeneity in human IFE has been largely unexplored compared to their murine counterparts. We made the surprising discovery of at least four basal stem cell populations in human neonatal epidermis using scRNA-seq and subsequent analysis with both Seurat and SoptSC. Each population is spatially distinct, with BAS-III cells occupying space between rete ridges and BAS-IV residing at the tips or bottom of the rete ridges, whereas the BAS-I and BAS-II populations showing sparse and heterogenous distribution throughout the basal and suprabasal layers. Analyzing cell-cell communication, gene profile dynamics, and genetic loss-of-function experiments indicate that the epigenetic modifiers HELLS and UHRF1 that reside in the BAS-II population, and the proto-oncogene PTTG1 that resides in the BAS-I population, are essential for epidermal homeostasis. Finally, our results provide clarity in the various models of IFE homeostasis and suggests that multiple stem cell pools with different proliferation capacities contribute to differentiation and epidermal homeostasis in humans.

The heterogeneity of basal stem cells in human IFE should not be surprising given that scRNA-seq studies have found robust heterogeneity in nearly all of the profiled tissues. However, KRT14 staining and use of the Krt14-Cre murine reporter lines as a proxy for the basal stem cell population lend to the inaccurate assumption of homogeneity in the epidermal basal layer. Use of mouse Cre reporter lines to untangle this homogeneity has led to models of IFE homeostasis that suggest two potentially distinct stem cell populations (Mascre et al., 2012; Sada et al., 2016), while others maintain one stem cell population (Clayton et al., 2007; Rompolas et al., 2016). Instead of the one or two basal stem cell populations from other models, our results indicate at least four spatially distinct basal populations exist in human neonatal epidermis. Intriguing hints in human studies point to this heterogeneity, where ITGB1 expression seemingly sits between rete ridges (Jensen et al., 1999), an observation we confirm with our ASS1 and COL17A1 immunostaining (Figure 2E). In fact, this population also appears to express *ITGA6* and *TP63*, suggesting it is the classically defined human basal stem cell population (Supplementary Figure 11). Additionally, ALP staining at the base of rete ridges (Lin et al., 2019) suggests another potentially unique basal population we confirm with our GJB2 immunostaining of the BAS-IV population (Figure 2G).

Our data indicates a linear hierarchy of differentiation beginning at the Krt14-CreER+/non-label retaining stem cell population (BAS-I), transitioning to the label retaining stem cell and *ITGB1*-high populations (BAS-II and BAS-III), and committing to differentiate in the Ivl-CreER+ committed progenitor population (BAS-IV). Once committed, the spinous cell populations appear to mature to terminally differentiated granular keratinocytes on a continuum where no gene clearly marks the spinous layer despite their respectively distinct morphologies. In practice, however, both the BAS-III and BAS-IV clusters appear to contribute to the spinous cell populations with RNA velocity vectors showing transitions from BAS-III to BAS-IV and SPN-1, whereas BAS-IV showing transitions to SPN-I, SPN-II, and GRN, further reinforcing a continuum of maturation from SPN-I to GRN. The spatial positioning of BAS-III and BAS-IV also suggest they both give rise to differentiated progeny as we observe KRT10 immunostaining beginning in the second layer along the rete ridges and flat portions of IFE (Figure 2D). The spatial positioning of the transitional basal populations (BAS-I and BAS-II), so-called because of their unusual spatial position where a significant percentage of the cells remain attached to the underlying basement membrane while their cell body is in a suprabasal position, suggests they may be delaminating from the basal layer. However, these basal cells are still attached to the basement membrane and may remain in the basal layer over time. Live imaging of these fate transitions would help shed light on their movements.

Why does interfollicular epidermal homeostasis need multiple stem cell populations? Skin appendages originate from the epidermal layer and it is possible that the epidermis actively responds to inductive signals even in the adult. For instance, normal epithelial cells are required for mutant epithelial cells resulting in aberrant growths to regress and are either eliminated or converted into functional skin appendages (Brown et al., 2017), suggesting that basal cell heterogeneity may be protective against harmful insults. In addition, basal stem cell self-renewal is not a constitutive driver of epidermal homeostasis; rather, cells need the plasticity to respond to their environment and coordinate divisions with stochastic loss of neighboring cells to maintain homeostasis (Mesa et al., 2018). At the very least, our results indicate that the transitional basal populations (BAS-I and BAS-II) are essential for epidermal homeostasis in human skin equivalents. Disruption of *PTTG1* expression (BAS-I) results in severe loss of epidermal stratification, resembling simple epithelium and displaying a more severe phenotype than was previously reported (Ishitsuka et al., 2012). And disruption of either *HELLS* or *UHRF1* expression (BAS-II) results in suppression of epidermal homeostasis and a thinner epidermis. HELLS is thought to recruit UHRF1 and DMNT1 to sites of methylated chromatin to facilitate demethylation and all three show strong similarities when disrupted in human skin equivalents (HELLS and UHRF1: Figure 4I; DMNT1: Sen et al., 2010), in stark contrast to the mouse phenotype where Krt14-Cre-mediated loss of *Dmnt1* results in an increase in IFE proliferation and aberrant differentiation (Li et al., 2012; JID). Differences in the frequency of KI67+ cells in *HELLS* and *UHRF1* KD skin equivalents suggest distinct differences between their functional roles in human epidermis despite their overlapping role in DNA demethylation.

Overall, our findings illustrate the dynamic nature of basal keratinocytes in human neonatal epidermis. The signaling and lineage relationships between the basal stem cell populations and differentiated keratinocytes warrants further characterization of the factors influencing fate plasticity and may help uncover the similarities and differences between vertebrate epidermal homeostasis and disease.

## MATERIALS AND METHODS

### Ethics statements

Human clinical studies were approved by the Ethics Committee of the University of California, Irvine. All human studies were performed in strict adherence to the Institutional Review Board (IRB) guidelines of the University of California, Irvine (2009-7083).

### Histology and immunohistochemistry

Frozen tissue sections (10μm) were fixed with 4% PFA in PBS for 15 minutes. Following fixation, tissue sections were stained with Hematoxylin and Eosin following standard procedures. Sections were stained with Gill’s III (Fisher Scientific; 22050203) for 5 minutes and Eosin-Y (Fisher Scientific; 22050197) for 1 minute. Tissue sections were visualized under a light microscope under 10x objective lens after mounting with Permount mounting media (Fisher Scientific; SP15-100). For immunostaining, tissue sections were fixed with 4% PFA in PBS for 15 minutes. 10% BSA in PBS was used for blocking. Following blocking, 5% BSA and 0.1% Triton X-100 in PBS was used for permeabilization. The following antibodies were used: chicken anti-KRT14 (1:500; BioLegend; SIG-3476), mouse anti-KRT10 (1:500; Dako; M7002), rabbit anti-KI67 (1:500; Abcam; ab15580), rabbit anti-PTTG1 (1:100; Sigma-Aldrich; HPA008890), mouse anti-ASS1 (1:100; Santa Cruz; sc-365475), rabbit anti-COL17A1 (1:100; One World Labs; ap9099c), rabbit anti-COL7A1 (1:500; abcam; ab93350), mouse anti-COL7A1 (1:100; Santa Cruz; sc-33710), rabbit anti-GJB2 (1:250; ThermoFisher; 51-2800), rabbit anti-KRT19 (1:250; Cell signaling; 13092), mouse anti-SPINK5 (1:100; Santa Cruz; sc-137109), mouse anti-CALML5 (1:100; Santa Cruz; sc-393637), mouse anti-CDC20 (1:100; Santa Cruz; sc-13162), mouse anti-RRM2 (1:100; Santa Cruz; sc-398294, sc-376973), mouse anti-PCLAF (1:100; Santa Cruz; sc-390515), mouse anti-MLANA (1:100; Santa Cruz; sc-20032), and mouse anti-CD74 (1:100; Santa Cruz; sc-6262). Secondary antibodies included Alexa Fluor 488 (1:500; Jackson ImmunoResearch; 715-545-150, 711-545-152) and Cy3 AffiniPure (1:500; Jackson ImmunoResearch; 711-165-152, 111-165-003). Slides were mounted with Prolong Diamond Antifade Mountant containing DAPI (Molecular Probes;). Confocal images were acquired at room temperature on a Zeiss LSM700 laser scanning microscope with Plan-Apochromat 20x objective or 40x and 63x oil immersion objectives. Images were arranged with ImageJ, Affinity Photo, and Affinity Designer.

### Protein immunoblotting

Cells were lysed with 2X SDS sample buffer (100 mM Tris HCl 6.8, 200 nM DTT, 4% SDS, 0.2% bromophenol blue, and 20% glycerol) and boiled at 100°C for 10 min. Samples were resolved on a 12.5% polyacrylamide gel and transferred to nitrocellulose membrane by a wet transfer apparatus. Membranes were blocked with 5% milk in TBS with 0.05% Tween-20 before sequential addition of primary and secondary antibodies. Membranes were imaged using Alexa Fluor secondary antibodies and the LI-COR Odyssey imaging system. Secondary antibodies included Alexa Fluor 680 (1:2000; Jackson ImmunoResearch; 715-625-150, 711-625-152) and Alexa Fluor 790 (1:2000; Jackson ImmunoResearch; 711-655-152)

### Lentiviral knockdown

Either pSicoR or pGIPZ vectors were used for lentiviral knockdown of specific genes. pSicoR was a gift from Tyler Jacks (Addgene; 11579). For pSicoR, shRNA were designed using pSicoligomaker 3.0 (Ventura Lab) and cloned using InFusion HD Cloning Plus Kit (Takara, 638911). The following sequences were used: PTTG1 5’- GATGATGCCTATCCAGAAATTCAAGAGATTTCTGGATAGGCATCATC-3’; RRM2 5’- GCACTCTAATGAAGCAATATTCAAGAGATATTGCTTCATTAGAGTGC-3’. Lentiviral pGIPZ vectors contained shRNAs to *UHRF1* (5′-TGACATTGCGCACCACCCT-3′) and *HELLS* (5′- ACAGGCTGATGTGTACTTAACC-3′; Wu and Benavente, 2018) (Dharmacon). Transduced cells were selected via Puromycin (ThermoFisher; 50464455). Knockdown efficiency was determined by protein levels on Western Blot or by quantitative RT-PCR where fold change in mRNA expression was measured using ΔΔC_T_ analysis with *GAPDH* as an internal control. The following qPCR primers were used: PTTG1 forward 5’-CCCTTGAGTGGAGTGCCTCT-3’, reverse 5’- CACAGCAAACAGGTGGCAAT-3’; RRM2 forward 5’-GCAGCAAGCGATGGCATAGT-3’, reverse 5’-GGGCTTCTGTAATCTGAACTTC-3’; PCNA forward 5’-CGACACCTACCGCTGCGACC-3’, reverse 5’-TAGCGCCAAGGTATCCGCGT-3’; and GAPDH forward 5’- GCACCGTCAAGGCTGAGAAC-3’, reverse 5’-TGGTGAAGACGCCAGTGGA-3’.

### Preparation of devitalized dermis

Cadaver human skin was acquired from the New York Firefighters Skin Bank (New York, New York, USA). Upon arrival at UC Irvine, the skin was allowed to thaw in a biosafety hood. Skin was then placed into PBS supplemented with 4X Pen/Strep, shaken vigorously for 5 minutes, and transferred to new PBS supplemented with 4X Pen/Strep. This step was repeated two additional times. The skin was then placed into a 37°C incubator for 2 weeks. The epidermis was removed from the dermis using using sterile watchmaker forceps. The dermis was washed 3 times in PBS supplemented with 4X Pen/Strep with vigorous shaking. The dermis was then stored in PBS supplemented with 4X Pen/Strep at 4°C until needed.

### Primary cell isolation

Discarded and de-identified neonatal foreskins were collected during routine circumcision from Orange County Medical Center (Santa Ana, CA, US). The samples were either processed for histological staining, single cell RNA-sequencing, or primary culture. No personal information was collected for this study. For primary cell isolation, fat from discarded and de-identified neonatal foreskins were removed using forceps and scissors and incubated with dispase epidermis side up for 2 hours at 37°C. The epidermis was peeled from the dermis, cut into fine pieces, and incubated in 0.25% Trypsin-EDTA for 15 minutes at 37°C and quenched with chelated FBS. Cells were passed through a 40µm filter, centrifuged at 1500rpm for 5 minutes, and the pellet resuspended in Keratinocyte Serum Free Media supplemented with Epidermal Growth Factor 1-53 and Bovine Pituitary Extract (Life Technologies; 17005042). Cells were either live/dead sorted using SYTOX Blue Dead Cell Stain (ThermoFisher; S34857) for single cell RNA-sequencing or incubated at 37°C for culture.

### Cell sorting

Following isolation, cells were resuspended in PBS free of Ca^2+^ and Mg^2+^ and 1% BSA and stained with SYTOX Blue Dead Cell Stain (ThermoFisher; S34857). Samples were bulk sorted at 4°C on a BD FACSAria Fusion using a 100µm nozzle (20 PSI) at a flow rate of 2.0 with a maximum threshold of 3000 events/sec. Following exclusion of debris and singlet/doublet discrimination, cells were gated on viability for downstream scRNA-seq.

### Human organotypic skin culture

Primary human keratinocytes were cultured in Keratinocyte Serum Free Media supplemented with Epidermal Growth Factor 1-53 and Bovine Pituitary Extract (Life Technologies; 17005042). For generating organotypic skin cultures, ∼500K control or knockdown cells were seeded on devitalized human dermis and raised to an air/liquid interface in order to induce differentiation and stratification as previously described (Li and Sen, 2015).

### Droplet-enabled single cell RNA-sequencing and processing

Cell counting, suspension, GEM generation, barcoding, post GEM-RT cleanup, cDNA amplification, library preparation, quality control, and sequencing was performed at the Genomics High Throughput Sequencing Facility at the University of California, Irvine. Transcripts were mapped to the human reference genome (GRCh38) using Cell Ranger Versions 1.3.0 and 2.1.0.

### Data repository

3’-end single cell RNA-sequencing data is in the process of submission to the GEO repository.

### Quality control metrics post-Cell Ranger assessment

For downstream analyses, we kept cells which met the following filtering criteria per biological replicate per condition: <6000 UMI/cell, and <10% mitochondrial gene expression. Genes that were expressed in less than 3 cells were excluded. Data was normalized with a scale factor of 10,000.

### Analysis and visualization of processed sequencing data

Seurat and SoptSC (Wang et al., 2019) were implemented for analysis of scRNA-seq data in this study. *<u>Seurat</u>.* (1.1) Implementation: Seurat was performed in R (version 2.2) and was applied to all the datasets in this study. (1.2) Unsupervised clustering and visualization: To select highly variable genes for initial clustering of cells, we performed Principal Component Analysis (PCA) on the scaled data for all genes included in the previous step. For clustering, we used the function *FindClusters* that implements Shared Nearest Neighbor (SNN) modularity optimization-based clustering algorithm on 20 PC components with resolution 0.6. Nonlinear dimensionality reduction methods, namely tSNE and UMAP, were applied to the scaled matrix for visualization of cells in two-dimensional space using first 10 PC components. (1.3) Gene selection: The marker genes for every cluster compared to all remaining cells were identified using the *FindAllMarkers* function. For each cluster, genes were selected such that they were expressed in at least 25% of cells with at least 0.25-fold difference. *<u>SoptSC</u>*. (2.1) Implementation: SoptSC was performed in MATLAB (version 2017b). In order to make both methods comparable for data clustering and downstream analysis, we used Seurat for quality control and normalization and used the resulting data matrices as an input for SoptSC. (2.2) Unsupervised clustering and visualization: Visualization of single cell data in low-dimensional space was implemented by applying Elastic Embedding (EE), a nonlinear dimensionality reduction technique, to the cell-to-cell similarity matrix. (2.3) Gene selection: SoptSC selected 3000 highly variable genes (HVGs) for the downstream analysis. The HVGs were identified via PCs on the single cell data. Genes with highest loadings in the first *k* principal components were selected, where the value of *k* is the index where the largest gap of the principal component variances occurs. The number of common marker genes between clusters among the top 500 marker genes for each cluster identified from two pipelines was compared to assess clustering correlations between Seurat and SoptSC.

### Pseudotime and lineage inference

Melanocytes and Langerhans cells were regressed out for lineage and pseudotime analysis. The resulting data post-melanocyte and Langerhans cell removal was re-clustered using SoptSC, where the number of HVGs was set to 10,000. Pseudotime and lineage analysis were performed using SoptSC. Briefly, pseudotime was calculated as the shortest path distance between cells and root cell on the cell-to-cell graph constructed based on the similarity matrix. Root cell was identified in an unsupervised manner in SoptSC. Visualization of the cell trajectories was obtained from EE with similarity matrix taken as an input. Cell states were visualized using abstract lineage trees. Lineage trees are obtained by computing the minimum spanning tree of the cluster-to-cluster graph based on the shortest path distance between cells. Pseudotime was projected on the lineage tree such that the order of each state (cluster) was defined as the average distances between cells within the state and the root cell.

### RNA velocity

RNA velocity was estimated based on the spliced and unspliced transcript reads from the single-cell data (La Manno et al., 2018). We followed the standard process of the velocyto pipeline to generate the spliced and unspliced matrices by applying *velocyto.py* to the data from the Cell Ranger output (outs) folder. We remove melanocytes and Langerhans cells for the velocity analysis and only epithelial cells included in pseudotime were used to calculate velocity vectors. RNA velocity was estimated using a gene-relative model with 25 nearest neighbors and then the velocity fields were projected onto the EE space produced by SoptSC. We set the parameter *n* sight as 2500, which defines the size of the neighborhood used for projecting velocity. We also applied the RNA velocity analysis for basal cells using the similar procedure where the parameter *n* was set as 500. Default settings were used for the rest of the parameters.

### Probabilistic cell-cell signaling networks

Cell-cell communication was determined in SoptSC via signaling networks for JAK/STAT, NOTCH, TGF-β, and WNT signaling pathways. In such networks, the probability between two cells is quantified by interactions between specific ligand-receptor pairs by the following equation as previously described (Wang et al., 2019). Prior calculation of the cell-cell interaction probability, we performed nature log normalization of the input count matrix. The lists of ligand-receptor pairs were determined before (Ramilowski et al., 2015; Subramanian et al., 2005) and with a survey of current literature. Circus plots (R Studio Version) are utilized as a visualization of cell-cell signaling networks. Edges between cells represent an interaction between them, and the width of each edge represents the interaction probability. Arrows start from ligand and points to receptor. We set a threshold (φ=0.1) such that the probability is restricted to zero if its value is less than φ. We display the top scoring 200 edges from the overall interactions for visualization due to the difficulty of circus plots with large edges. Cluster-level communications were naturally employed by calculating the average value of probabilities between cell-cell interactions from two clusters. In the cluster-to-cluster signaling network plots for each specific pathway, the width of the edge represents the probability value between clusters. Arrows start from ligand and points to receptor. Ligand-receptor pairs and their targets genes among all inferred cell populations are also presented as heatmaps. For each gene, the average expression within each population was calculated.

### Gene Ontology Analysis

Top 100 differentially expressed genes from each basal cluster were used for gene ontology and pathway analyses using Enrichr (Chen et al., 2013; Kuleshov et al., 2016).

### Cellular entropy estimation

Cellular Entropy (*ξ*) measures the likelihood that a cell will transition to a new state (i.e. from one cluster to another). Lower entropy values indicate that the cell remains in a steady state, while higher entropy values imply the cell inherits multiple state properties and is more likely to transition to a new state. Via the NMF step in SoptSC, the probability of each cell assigned to each cluster is calculated (i.e., *P_i,j_* for cell *i* and cluster *j*). The entropy for each cell is then defined as:

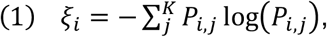

where *K* represents the number of clusters. To visualize the trajectory of cells and their likelihood of transition along a transition valley, we constructed a Waddington landscape and overlaid the cell states on it. The Waddington landscape is constructed by integrating the low-dimensional representation of our data (via applying EE to similarity matrix) and the cellular entropy estimation for each cell. This new feature has been extended to the current methods employed by SoptSC.

## ACKNOWLEDGEMENTS

S.X.A. is supported by the Concern Foundation (CF204525) and start-up funds from UCI. Q.N. is supported by NIH grants R01GM123731 and U01AR07315, a NSF grants DMS11763272, a grant by Jayne Koskinas Ted Giovanis Foundation for Health and Policy jointly with the Breast Cancer Research Foundation, and a NSF-Simons Foundation grant (594598). C.F.G.-J. is supported by UC Irvine Chancellor’s ADVANCE Postdoctoral Fellowship, NSF-Simons Postdoctoral Fellowship, and a NSF Grant DMS11763272. E.T. and A.R.S. are supported by the GAANN Fellowship. This project was partly funded by an opportunity award from the UCI Center for Complex Biological Systems (P50GM076516) and the UCI Office of Research. We thank Jennifer Bates and the Institute for Immunology Flow Cytometry Core Facility for help with cell sorting.

## CONTRIBUTIONS

S.X.A. and Q.N. conceived the project; S.X.A. and Q.N. supervised research; M.L.D. generated scRNA-seq libraries; S.W. and M.L.D. analyzed scRNA-seq data; M.L.D., E.T., G.G., and A.R.S. performed imaging experiments; M.L.D., E.T., S.C.W., and B.T.T. performed knockdown experiments; S.W., C.F.G.-J., A.L.M., M.L.D., Q.N., and S.X.A. analyzed and interpreted data; C.A.B. contributed *HELLS* and *UHRF1* KD data and reagents; S.X.A., C.F.G.-J., and S.W. wrote the manuscript. All authors analyzed and discussed the results and commented on the manuscript.

---

**Supplementary Figure 1.**
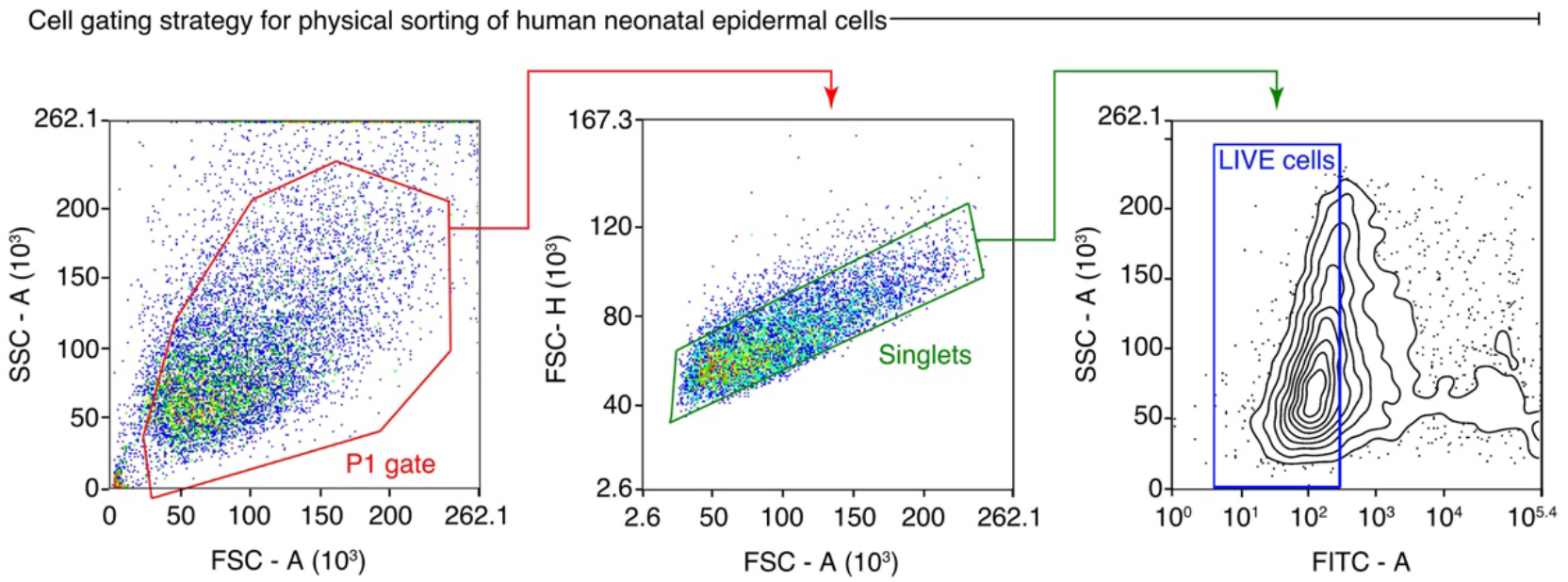
Schematic of FACS strategy for physical sorting of live epidermal cells from human neonatal epidermis samples.

**Supplementary Figure 2.**
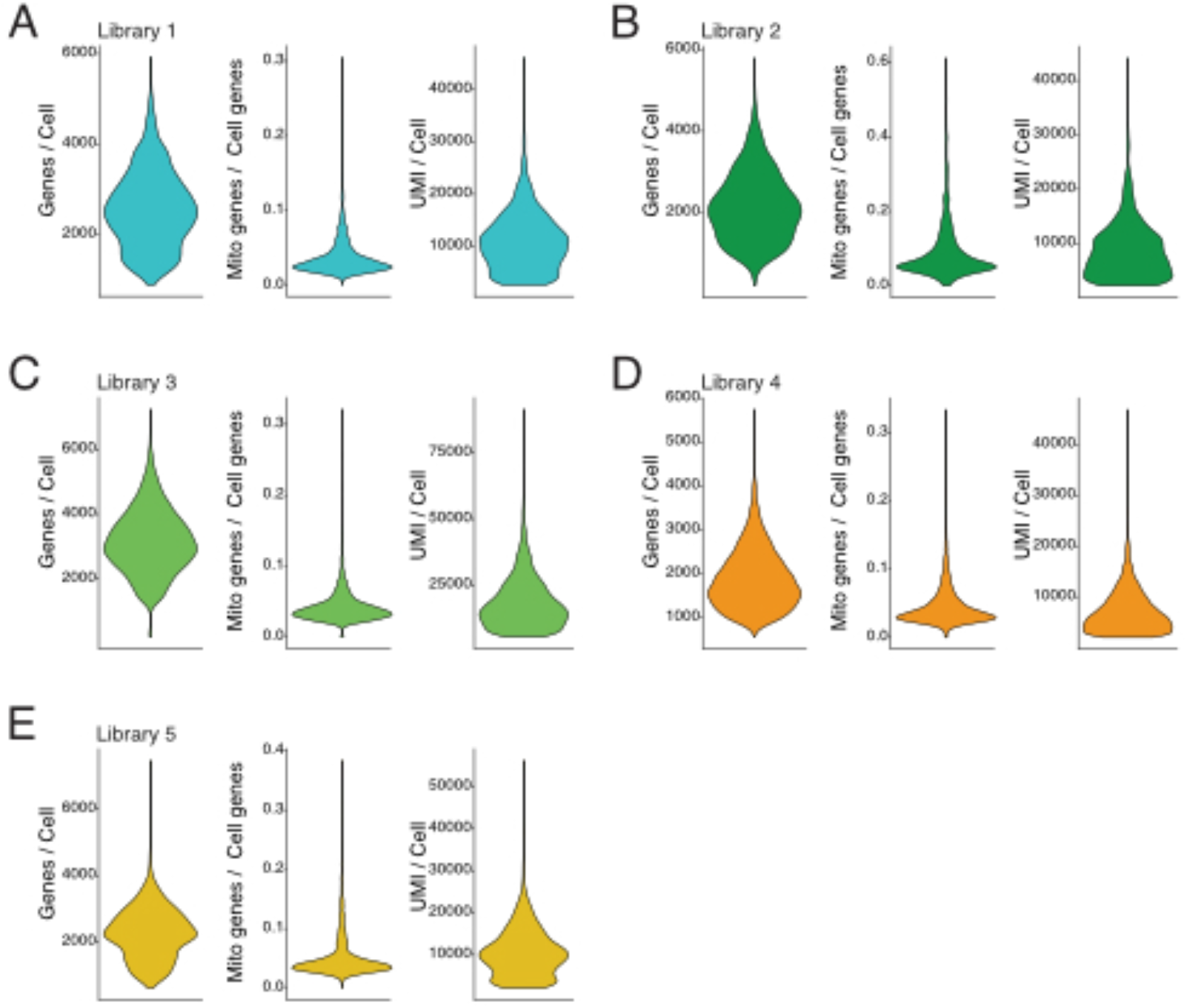
Metrics of single cell libraries. **A-E)** Violin plots showing genes per cell, mitochondrial (mito) genes per total cell genes, and unique molecular identifiers (UMI) per cell during quality control assessment.

**Supplementary Figure 3.**
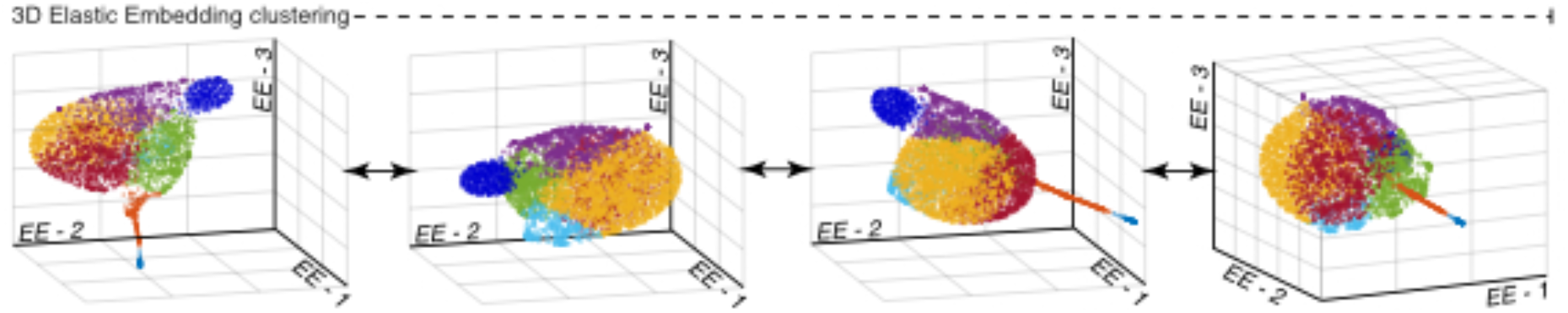
3D clustering of epidermal cells. Three-dimensional (3D) elastic embedding (EE) clustering of epidermal cells isolated from human neonatal epidermis. Cells color-coded as in Figure 1B.

**Supplementary Figure 4.**
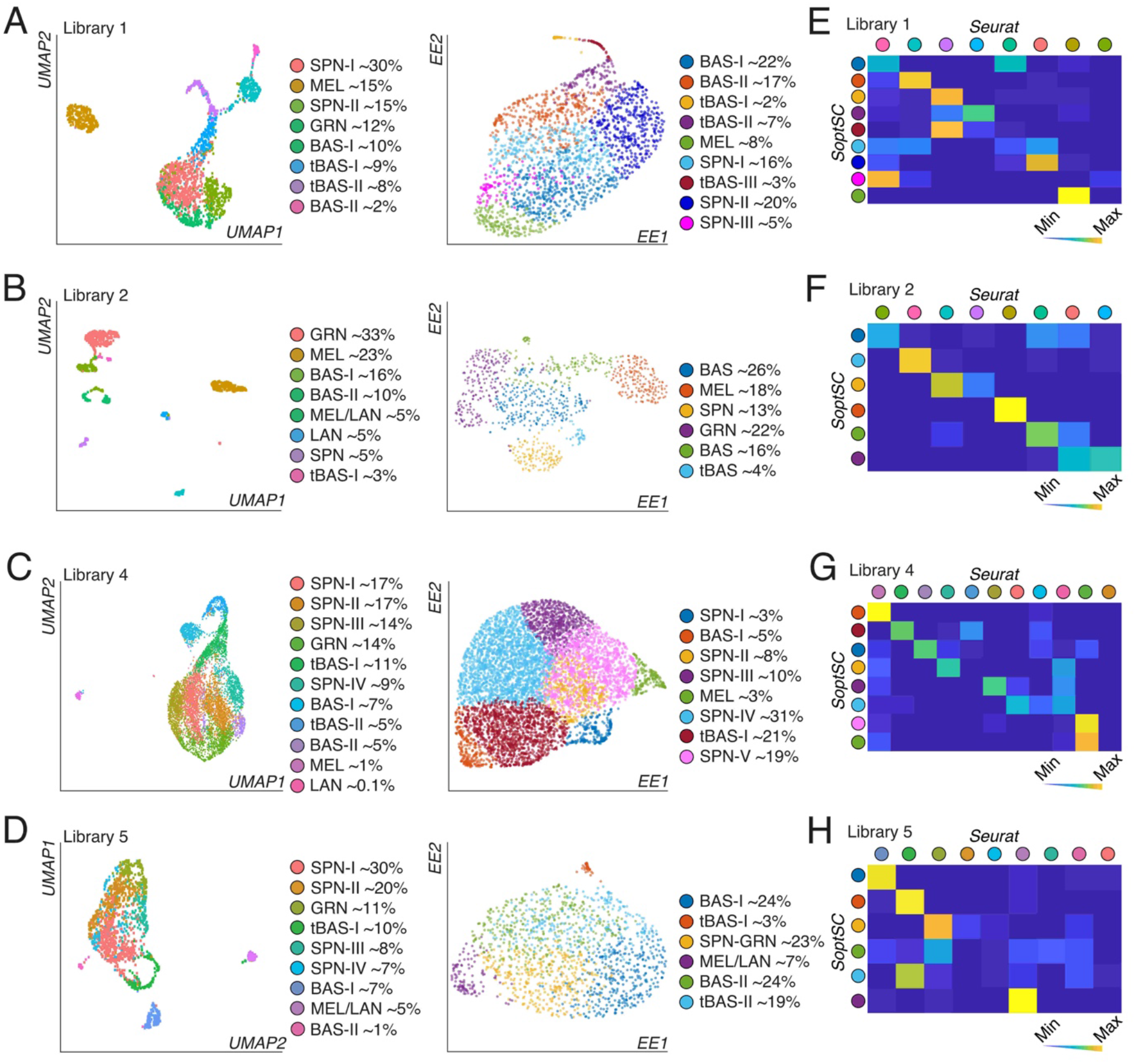
Clustering and correlation analysis of epidermal cells. **A-D)** Clustering of single cells isolated from human neonatal epidermis using (left) Seurat and dimensionality reduction using UMAP and (right) using SoptSC and displayed using EE. Cell proportions from putative cellular communities are quantified on the right of each plot. Library numbers as labeled. **E-H)** Correlation between overlapping differentially expressed genes from SoptSC and Seurat. Library numbers as labeled.

**Supplementary Figure 5.**
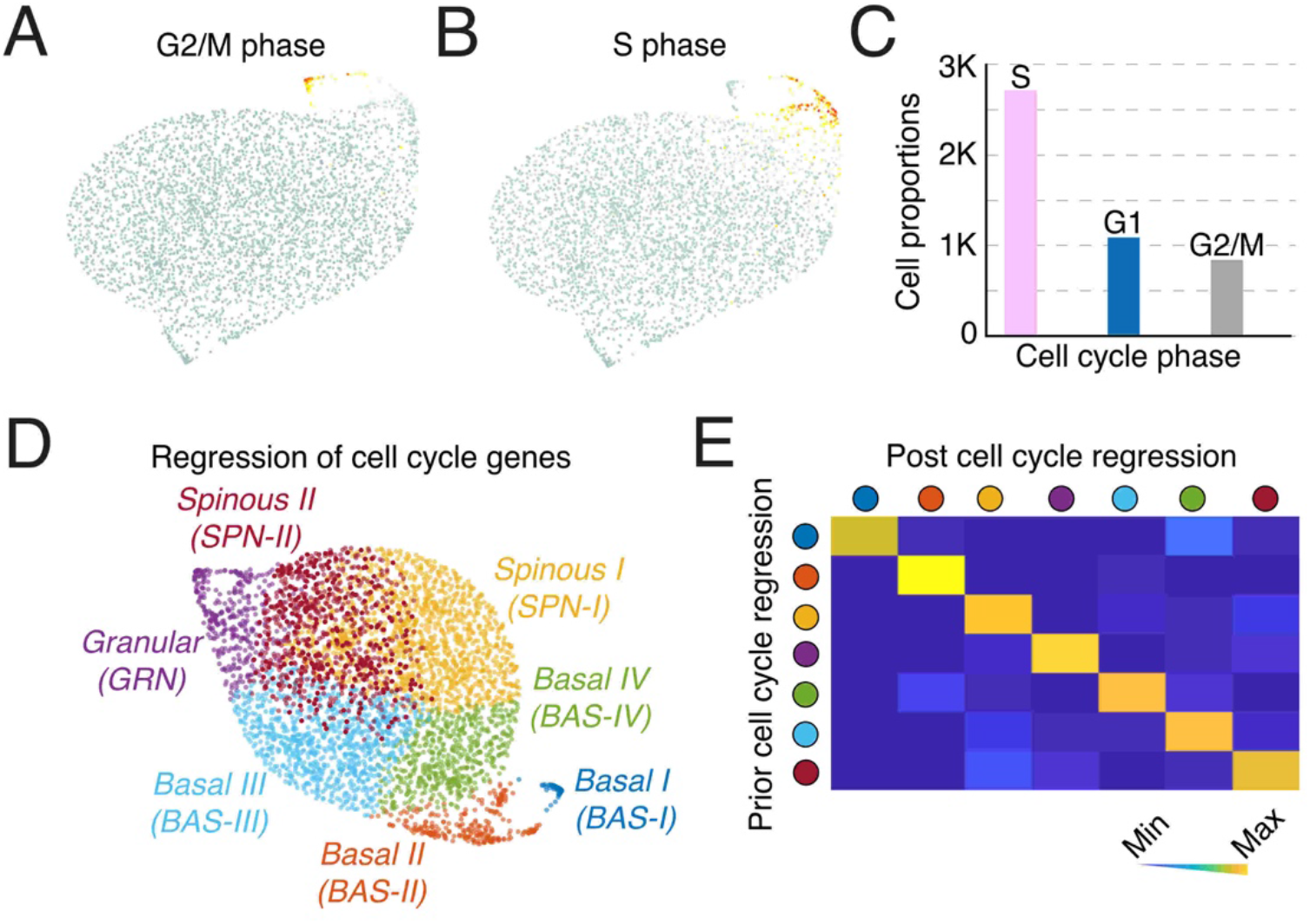
Cell cycle analysis of epidermal cells. **A)** G2/M phase score overlaid on EE space of Library 3. **B)** S phase score overlaid on EE space. **C)** Quantification of cell cycle phase state. K, 1000. **D)** Clustering of epidermal cells after regression of cell cycle genes. **E)** Correlation between overlapping markers before and after cell cycle regression.

**Supplementary Figure 6.**
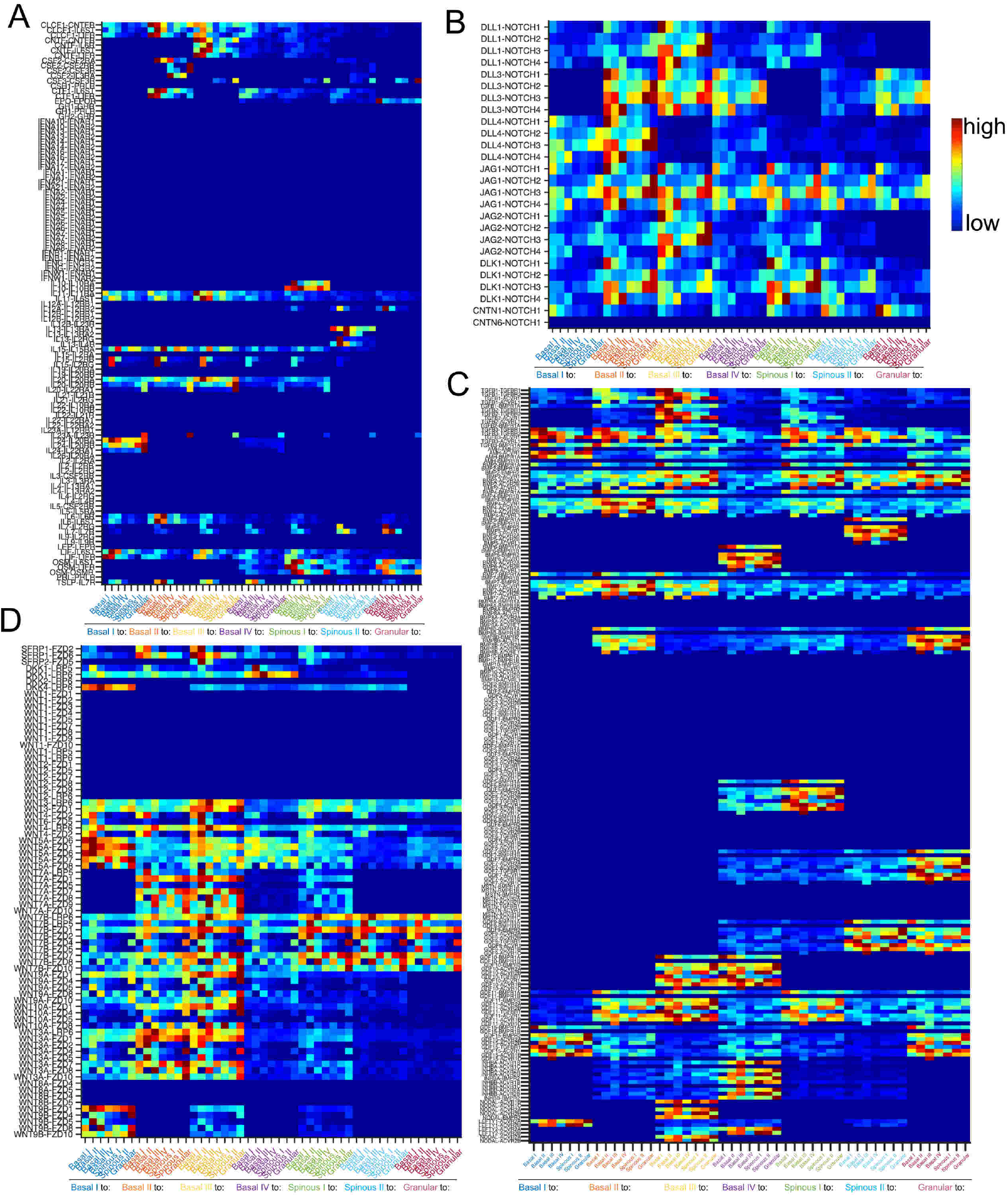
Cell-cell communication heatmaps. **A-D)** Cell-cell communication heatmaps predicted for the (A) JAK-STAT, (B) NOTCH, (C) TGF-β, and (D) WNT signaling pathways. Left: Ligand-receptor pairs (downstream targets for each pair not labeled). Bottom: Cluster signaling relationships. Colors correspond to cluster labels from Figure 4A.

**Supplementary Figure 7.**
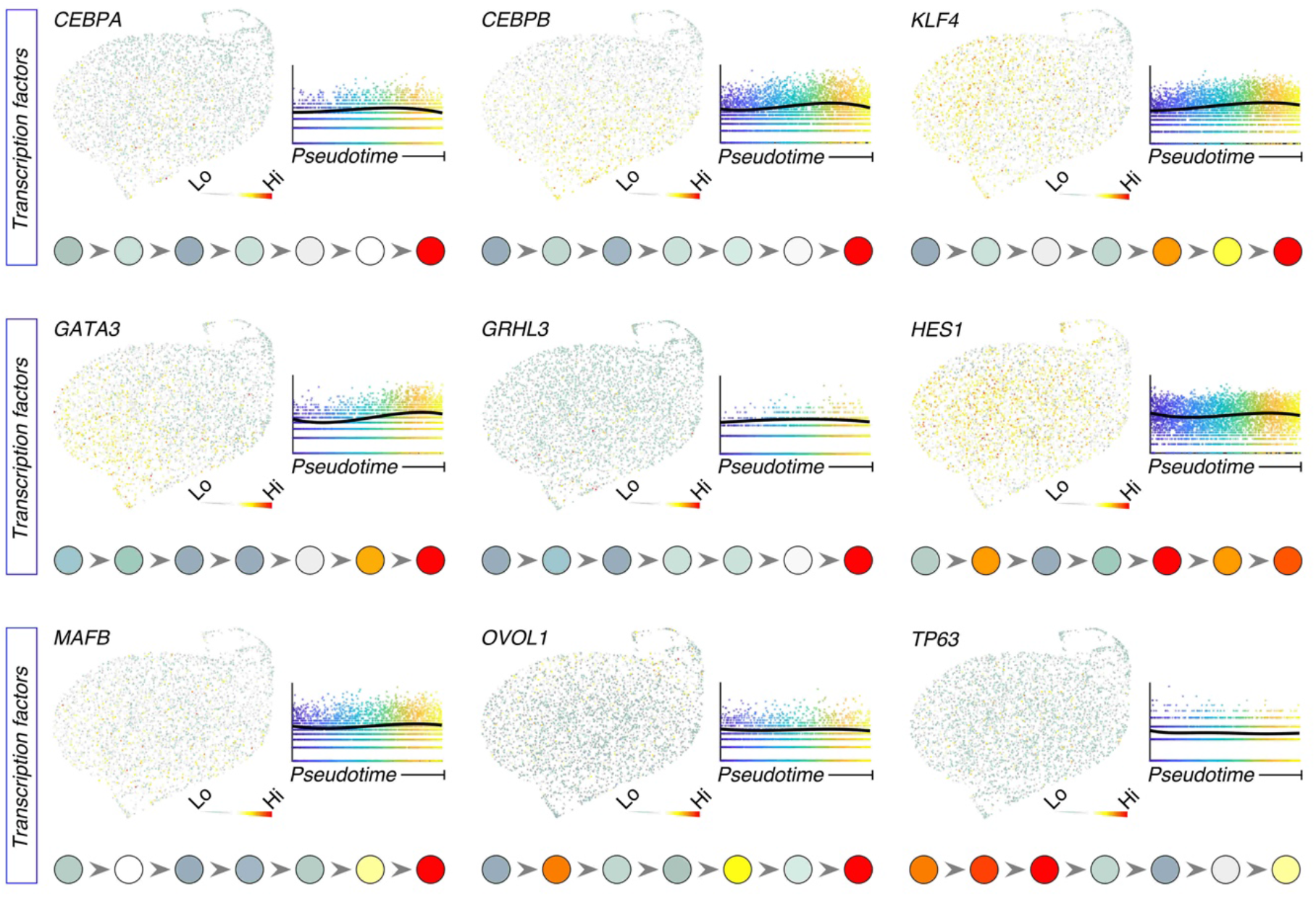
Evaluation of transcription factor dynamics along pseudotime. Feature plots showing expression of select transcription factor genes and their expression along pseudotime. Expression along the epidermal cell lineage as determined in Figure 4B displayed at the bottom.

**Supplementary Figure 8.**
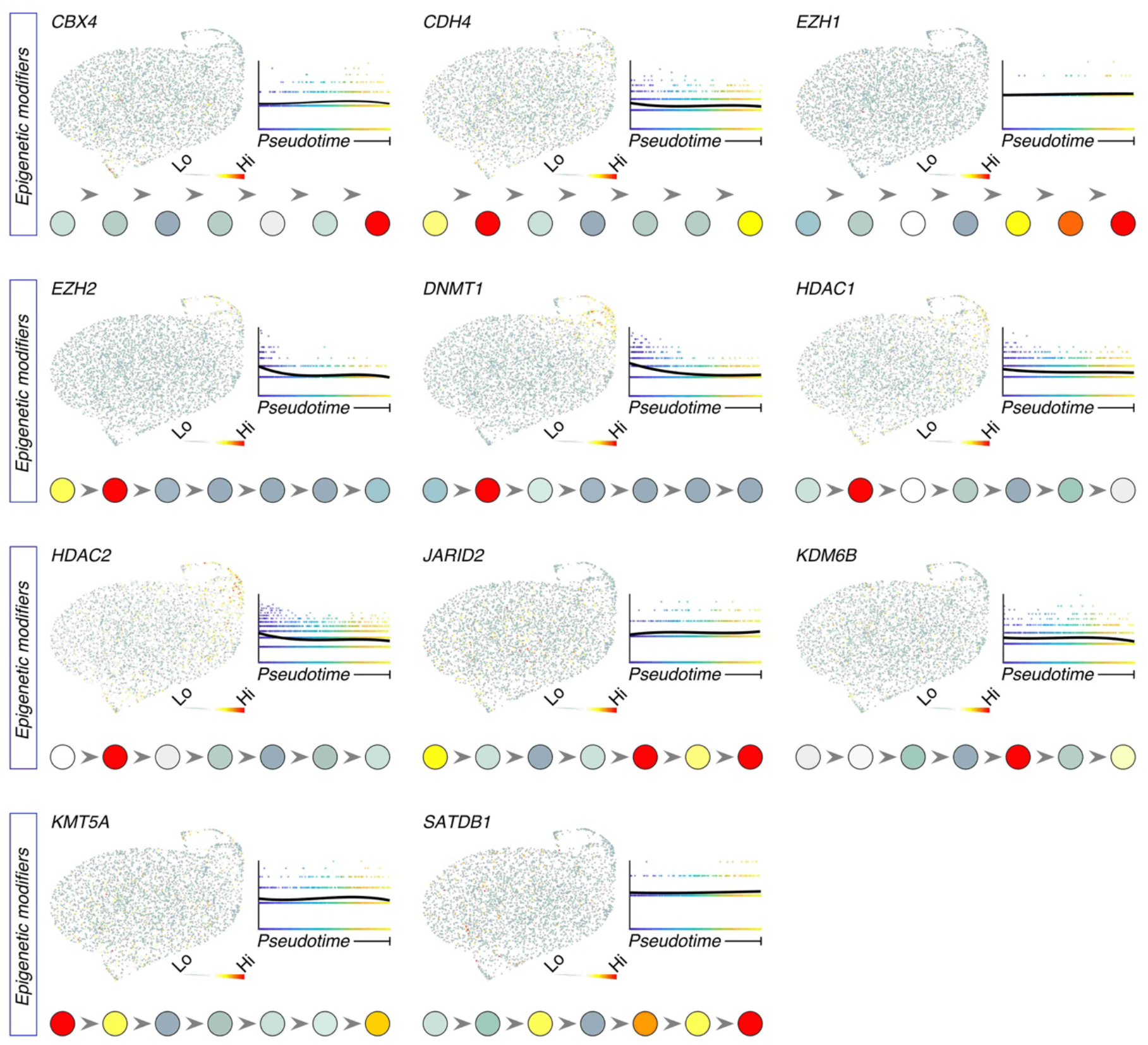
Evaluation of epigenetic modifier dynamics along pseudotime. Feature plots showing expression of select epigenetic modifier genes and their expression along pseudotime. Expression along the epidermal cell lineage as determined in Figure 4B displayed at the bottom.

**Supplementary Figure 9.**
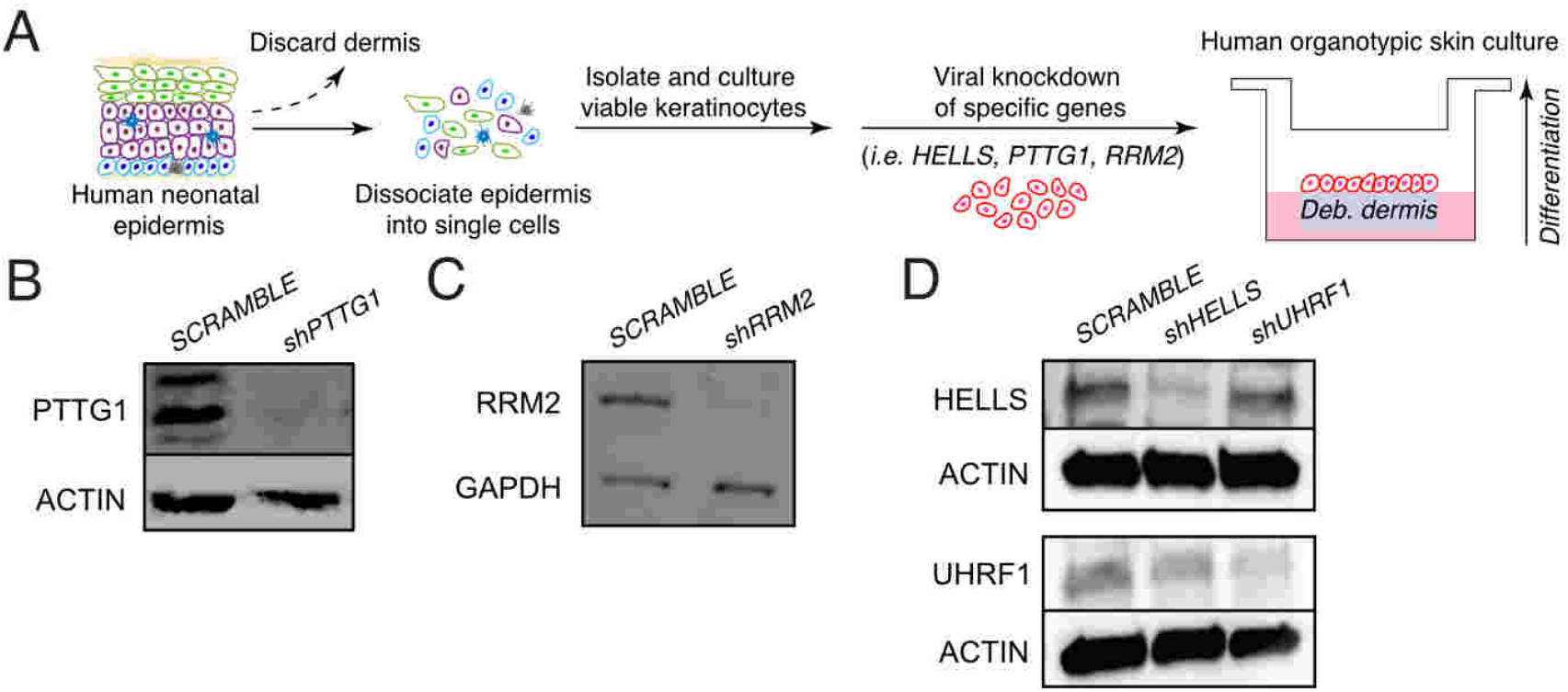
Generation of human organotypic skin cultures. **A)** Schematic of primary keratinocyte cell isolation from human neonatal epidermis, viral knockdown, and seeding on top of devitalized human dermis at the air-liquid interface for generation of organotypic skin cultures. **B)** Western blot of PTTG1 or ACTB (ACTIN) protein in *SCRAMBLE* control or *PPTG1* knockdown (KD) (*shPTTG1*) keratinocytes. **C)** Western blot of RRM2 or GAPDH protein in *SCRAMBLE* control or *RRM2* KD (*shRRM2*) keratinocytes. **D)** Western blot of HELLS, UHRF1, or ACTIN protein in *SCRAMBLE* control, *HELLS* KD (*shHELLS*), or *UHRF1* KD (*shUHRF1*) keratinocytes.

**Supplementary Figure 10.**
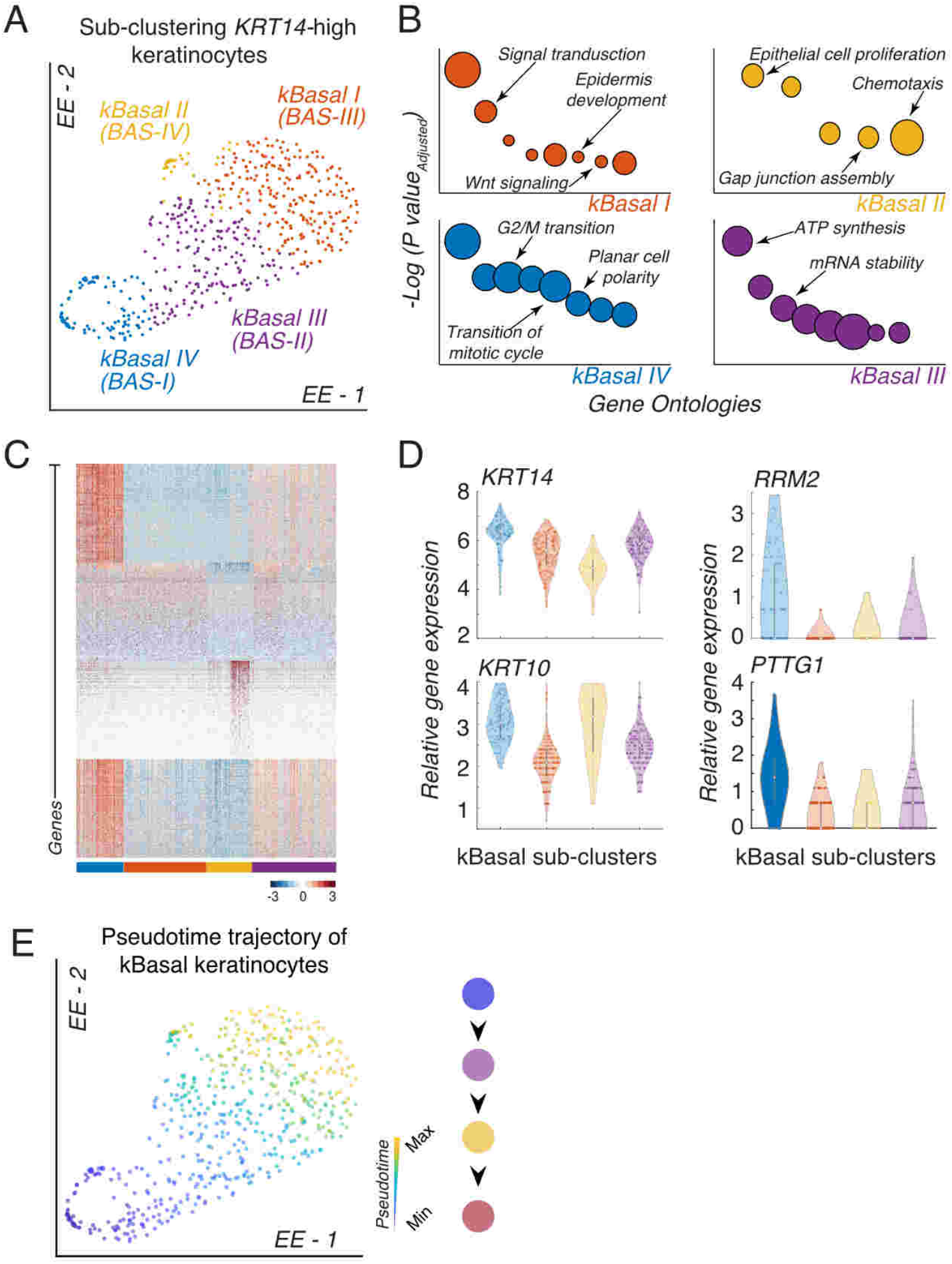
Subclustering KRT14-high cells subsample existing basal stem cell populations with no increase in heterogeneity. **A)** Subclustering of *KRT14*-high keratinocytes using SoptSC and displayed using EE. Corresponding cluster identification from Figure 4A in parenthesis. **B)** Gene ontology analysis of each kBasal cluster with selected terms labeled. **C)** Heatmap showing top 100 differentially expressed genes per cluster. Cells are color-coded as in A at the bottom. **D)** Violin plot of relative gene expression of *KRT14*, *KRT10*, *PTTG1*, and *RRM2* across the kBasal subclusters. **E)** Pseudotime inference of kbasal cluster keratinocytes displayed using EE. Cell lineage inference displayed on the right.

**Supplementary Figure 11.**
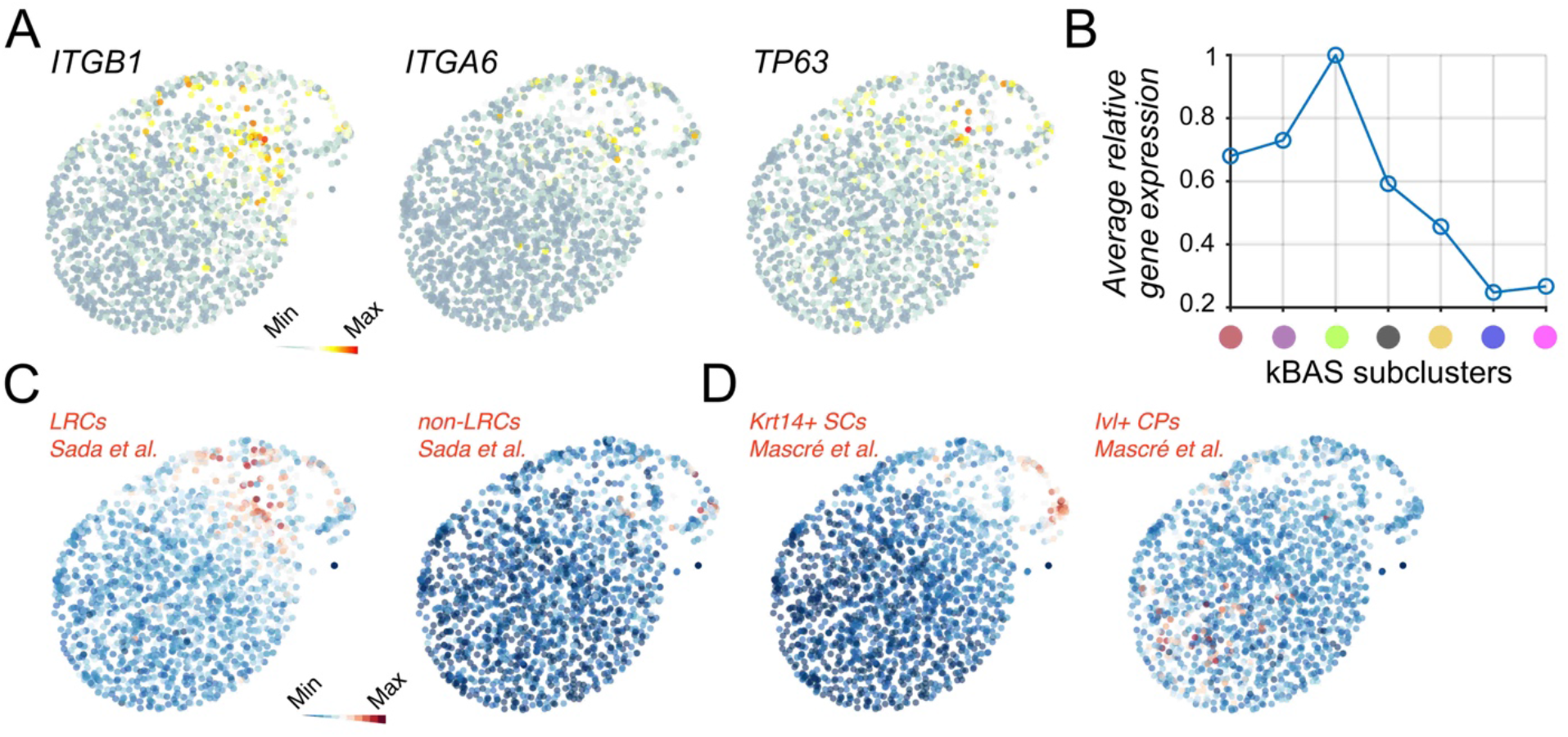
Characterization of markers associated with stem cells and committed progenitors. **A)** Feature plots showing expression of common stem cell-associated genes (*ITGB1*, *ITGA6*, and *TP63*) overlaid on EE space. **B)** Average relative gene expression of stem cell-associated genes along the kBAS subclusters from Figure 5A. **C)** Feature plots showing average relative gene expression of label-retaining stem cells (LRC) or non-label-retaining stem cells (non-LRC) from Sada et al., 2016. D) Feature plots showing average relative gene expression of Krt14-CreER+ (Krt14+) stem cells (SC) or Ivl-CreER+ (Ivl+) committed progenitors (CP) from Mascre et al., 2012.

## REFERENCES

Aird, K.M., Zhang, G., Li, H., Tu, Z., Bitler, B.G., Garipov, A., Wu, H., Wei, Z., Wagner, S.N., Herlyn, M., et al. (2013). Suppression of nucleotide metabolism underlies the establishment and maintenance of oncogene-induced senescence. Cell Rep 3, 1252–1265.

Andl, T., Reddy, S.T., Gaddapara, T., and Millar, S.E. (2002). WNT signals are required for the initiation of hair follicle development. Dev Cell 2, 643–653.

Bamberger, C., Scharer, A., Antsiferova, M., Tychsen, B., Pankow, S., Muller, M., Rulicke, T., Paus, R., and Werner, S. (2005). Activin controls skin morphogenesis and wound repair predominantly via stromal cells and in a concentration-dependent manner via keratinocytes. Am J Pathol 167, 733–747.

Benavente, C.A., Finkelstein, D., Johnson, D.A., Marine, J.C., Ashery-Padan, R., and Dyer, M.A. (2014). Chromatin remodelers HELLS and UHRF1 mediate the epigenetic deregulation of genes that drive retinoblastoma tumor progression. Oncotarget 5, 9594–9608.

Bikle, D.D., Xie, Z., and Tu, C.L. (2012). Calcium regulation of keratinocyte differentiation. Expert Rev Endocrinol Metab 7, 461–472.

Blanpain, C., Lowry, W.E., Geoghegan, A., Polak, L., and Fuchs, E. (2004). Self-renewal, multipotency, and the existence of two cell populations within an epithelial stem cell niche. Cell 118, 635–648.

Blanpain, C., Lowry, W.E., Pasolli, H.A., and Fuchs, E. (2006). Canonical notch signaling functions as a commitment switch in the epidermal lineage. Genes Dev 20, 3022–3035.

Botchkarev, V.A., Gdula, M.R., Mardaryev, A.N., Sharov, A.A., and Fessing, M.Y. (2012). Epigenetic regulation of gene expression in keratinocytes. J Invest Dermatol 132, 2505–2521.

Brown, S., Pineda, C.M., Xin, T., Boucher, J., Suozzi, K.C., Park, S., Matte-Martone, C., Gonzalez, D.G., Rytlewski, J., Beronja, S., and Greco, V. (2017). Correction of aberrant growth preserves tissue homeostasis. Nature 548, 334–337.

Carreira-Perpiñan (2010). The elastic embedding algorithm for dimensionality reduction. In Proceedings of the 27th International Conference on International Conference on Machine Learning (Haifa, Israel: Omnipress), pp. 167–174.

Chen, E.Y., Tan, C.M., Kou, Y., Duan, Q., Wang, Z., Meirelles, G.V., Clark, N.R., and Ma’ayan, A. (2013). Enrichr: interactive and collaborative HTML5 gene list enrichment analysis tool. BMC Bioinformatics 14, 128.

Cheng, J.B., Sedgewick, A.J., Finnegan, A.I., Harirchian, P., Lee, J., Kwon, S., Fassett, M.S., Golovato, J., Gray, M., Ghadially, R., et al. (2018). Transcriptional Programming of Normal and Inflamed Human Epidermis at Single-Cell Resolution. Cell Rep 25, 871–883.

Choi, Y.S., Zhang, Y., Xu, M., Yang, Y., Ito, M., Peng, T., Cui, Z., Nagy, A., Hadjantonakis, A.K., Lang, R.A., et al. (2013). Distinct functions for Wnt/beta-catenin in hair follicle stem cell proliferation and survival and interfollicular epidermal homeostasis. Cell Stem Cell 13, 720–733.

Clayton, E., Doupe, D.P., Klein, A.M., Winton, D.J., Simons, B.D., and Jones, P.H. (2007). A single type of progenitor cell maintains normal epidermis. Nature 446, 185–189.

Clevers, H. (2006). Wnt/beta-catenin signaling in development and disease. Cell 127, 469–480.

Clevers, H., and Nusse, R. (2012). Wnt/beta-catenin signaling and disease. Cell 149, 1192–1205.

Cotsarelis, G., Sun, T.T., and Lavker, R.M. (1990). Label-retaining cells reside in the bulge area of pilosebaceous unit: implications for follicular stem cells, hair cycle, and skin carcinogenesis. Cell 61, 1329–1337.

Cottle, D.L., Kretzschmar, K., Schweiger, P.J., Quist, S.R., Gollnick, H.P., Natsuga, K., Aoyagi, S., and Watt, F.M. (2013). c-MYC-induced sebaceous gland differentiation is controlled by an androgen receptor/p53 axis. Cell Rep 3, 427–441.

de Guzman Strong, C., Wertz, P.W., Wang, C., Yang, F., Meltzer, P.S., Andl, T., Millar, S.E., Ho, I.C., Pai, S.Y., and Segre, J.A. (2006). Lipid defect underlies selective skin barrier impairment of an epidermal-specific deletion of Gata-3. J Cell Biol 175, 661–670.

Dong, J., Hu, Y., Fan, X., Wu, X., Mao, Y., Hu, B., Guo, H., Wen, L., and Tang, F. (2018). Single-cell RNA-seq analysis unveils a prevalent epithelial/mesenchymal hybrid state during mouse organogenesis. Genome Biol 19, 31.

Eckert, R.L., and Rorke, E.A. (1989). Molecular biology of keratinocyte differentiation. Environ Health Perspect 80, 109–116.

Ezhkova, E., Pasolli, H.A., Parker, J.S., Stokes, N., Su, I.H., Hannon, G., Tarakhovsky, A., and Fuchs, E. (2009). Ezh2 orchestrates gene expression for the stepwise differentiation of tissue-specific stem cells. Cell 136, 1122–1135.

Fessing, M.Y., Mardaryev, A.N., Gdula, M.R., Sharov, A.A., Sharova, T.Y., Rapisarda, V., Gordon, K.B., Smorodchenko, A.D., Poterlowicz, K., Ferone, G., et al. (2011). p63 regulates Satb1 to control tissue-specific chromatin remodeling during development of the epidermis. J Cell Biol 194, 825–839.

Geiman, T.M., Tessarollo, L., Anver, M.R., Kopp, J.B., Ward, J.M., and Muegge, K. (2001). Lsh, a SNF2 family member, is required for normal murine development. Biochim Biophys Acta 1526, 211–220.

Ghazizadeh, S., Katz, A.B., Harrington, R., and Taichman, L.B. (2004). Lentivirus-mediated gene transfer to human epidermis. J Investig Dermatol Symp Proc 9, 269–275.

Ghazizadeh, S., and Taichman, L.B. (2005). Organization of stem cells and their progeny in human epidermis. J Invest Dermatol 124, 367–372.

Greco, V., Chen, T., Rendl, M., Schober, M., Pasolli, H.A., Stokes, N., Dela Cruz-Racelis, J., and Fuchs, E. (2009). A two-step mechanism for stem cell activation during hair regeneration. Cell Stem Cell 4, 155–169.

Guerrero-Juarez, C.F., Dedhia, P.H., Jin, S., Ruiz-Vega, R., Ma, D., Liu, Y., Yamaga, K., Shestova, O., Gay, D.L., Yang, Z., et al. (2019). Single-cell analysis reveals fibroblast heterogeneity and myeloid-derived adipocyte progenitors in murine skin wounds. Nat Commun 10, 650.

Gupta, K., Levinsohn, J., Linderman, G., Chen, D., Sun, T.Y., Dong, D., Taketo, M.M., Bosenberg, M., Kluger, Y., Choate, K., et al. (2019). Single-Cell Analysis Reveals a Hair Follicle Dermal Niche Molecular Differentiation Trajectory that Begins Prior to Morphogenesis. Dev Cell 48, 17–31 e16.

Hanel, K.H., Cornelissen, C., Luscher, B., and Baron, J.M. (2013). Cytokines and the skin barrier. Int J Mol Sci 14, 6720–6745.

Heldin, C.H., and Moustakas, A. (2016). Signaling Receptors for TGF-beta Family Members. Cold Spring Harb Perspect Biol 8.

Horsley, V., O’Carroll, D., Tooze, R., Ohinata, Y., Saitou, M., Obukhanych, T., Nussenzweig, M., Tarakhovsky, A., and Fuchs, E. (2006). Blimp1 defines a progenitor population that governs cellular input to the sebaceous gland. Cell 126, 597–609.

Hughes, T.K., Wadsworth, M.H., Gierahn, T.M., Do, T., Weiss, D., Andrade, P.R., Ma, F., de Andrade Silva, B.J., Shao, S., Tsoi, L.C., et al. (2019). Highly Efficient, Massively-Parallel Single-Cell RNA-Seq Reveals Cellular States and Molecular Features of Human Skin Pathology. bioRxiv, 689273.

Indra, A.K., Dupe, V., Bornert, J.M., Messaddeq, N., Yaniv, M., Mark, M., Chambon, P., and Metzger, D. (2005). Temporally controlled targeted somatic mutagenesis in embryonic surface ectoderm and fetal epidermal keratinocytes unveils two distinct developmental functions of BRG1 in limb morphogenesis and skin barrier formation. Development 132, 4533–4544.

Ishitsuka, Y., Kawachi, Y., Maruyama, H., Taguchi, S., Fujisawa, Y., Furuta, J., Nakamura, Y., Ishii, Y., and Otsuka, F. (2013). Pituitary tumor transforming gene 1 induces tumor necrosis factor-alpha production from keratinocytes: implication for involvement in the pathophysiology of psoriasis. J Invest Dermatol 133, 2566–2575.

Ishitsuka, Y., Kawachi, Y., Taguchi, S., Maruyama, H., Fujisawa, Y., Furuta, J., Nakamura, Y., and Otsuka, F. (2012). Pituitary tumor-transforming gene 1 enhances proliferation and suppresses early differentiation of keratinocytes. J Invest Dermatol 132, 1775–1784.

Ito, M., and Kizawa, K. (2001). Expression of calcium-binding S100 proteins A4 and A6 in regions of the epithelial sac associated with the onset of hair follicle regeneration. J Invest Dermatol 116, 956–963.

Ito, M., Kizawa, K., Toyoda, M., and Morohashi, M. (2002). Label-retaining cells in the bulge region are directed to cell death after plucking, followed by healing from the surviving hair germ. J Invest Dermatol 119, 1310–1316.

Jensen, K.B., Collins, C.A., Nascimento, E., Tan, D.W., Frye, M., Itami, S., and Watt, F.M. (2009). Lrig1 expression defines a distinct multipotent stem cell population in mammalian epidermis. Cell Stem Cell 4, 427–439.

Jensen, U.B., Lowell, S., and Watt, F.M. (1999). The spatial relationship between stem cells and their progeny in the basal layer of human epidermis: a new view based on whole-mount labelling and lineage analysis. Development 126, 2409–2418.

Jensen, U.B., Yan, X., Triel, C., Woo, S.H., Christensen, R., and Owens, D.M. (2008). A distinct population of clonogenic and multipotent murine follicular keratinocytes residing in the upper isthmus. J Cell Sci 121, 609–617.

Jones, P.H., Simons, B.D., and Watt, F.M. (2007). Sic transit gloria: farewell to the epidermal transit amplifying cell? Cell Stem Cell 1, 371–381.

Joost, S., Jacob, T., Sun, X., Annusver, K., La Manno, G., Sur, I., and Kasper, M. (2018). Single-Cell Transcriptomics of Traced Epidermal and Hair Follicle Stem Cells Reveals Rapid Adaptations during Wound Healing. Cell Rep 25, 585–597 e587.

Joost, S., Zeisel, A., Jacob, T., Sun, X., La Manno, G., Lonnerberg, P., Linnarsson, S., and Kasper, M. (2016). Single-Cell Transcriptomics Reveals that Differentiation and Spatial Signatures Shape Epidermal and Hair Follicle Heterogeneity. Cell Syst 3, 221–237 e229.

Jung, H.J., Byun, H.O., Jee, B.A., Min, S., Jeoun, U.W., Lee, Y.K., Seo, Y., Woo, H.G., and Yoon, G. (2017). The Ubiquitin-like with PHD and Ring Finger Domains 1 (UHRF1)/DNA Methyltransferase 1 (DNMT1) Axis Is a Primary Regulator of Cell Senescence. J Biol Chem 292, 3729–3739.

Karaayvaz, M., Cristea, S., Gillespie, S.M., Patel, A.P., Mylvaganam, R., Luo, C.C., Specht, M.C., Bernstein, B.E., Michor, F., and Ellisen, L.W. (2018). Unravelling subclonal heterogeneity and aggressive disease states in TNBC through single-cell RNA-seq. Nat Commun 9, 3588.

Kashiwagi, M., Morgan, B.A., and Georgopoulos, K. (2007). The chromatin remodeler Mi-2beta is required for establishment of the basal epidermis and normal differentiation of its progeny. Development 134, 1571–1582.

Keyes, W.M., Pecoraro, M., Aranda, V., Vernersson-Lindahl, E., Li, W., Vogel, H., Guo, X., Garcia, E.L., Michurina, T.V., Enikolopov, G., et al. (2011). DeltaNp63alpha is an oncogene that targets chromatin remodeler Lsh to drive skin stem cell proliferation and tumorigenesis. Cell Stem Cell 8, 164–176.

Kratochwil, K., Dull, M., Farinas, I., Galceran, J., and Grosschedl, R. (1996). Lef1 expression is activated by BMP-4 and regulates inductive tissue interactions in tooth and hair development. Genes Dev 10, 1382–1394.

Kuleshov, M.V., Jones, M.R., Rouillard, A.D., Fernandez, N.F., Duan, Q., Wang, Z., Koplev, S., Jenkins, S.L., Jagodnik, K.M., Lachmann, A., et al. (2016). Enrichr: a comprehensive gene set enrichment analysis web server 2016 update. Nucleic Acids Res 44, W90–97.

Kumar, M.P., Du, J., Lagoudas, G., Jiao, Y., Sawyer, A., Drummond, D.C., Lauffenburger, D.A., and Raue, A. (2018). Analysis of Single-Cell RNA-Seq Identifies Cell-Cell Communication Associated with Tumor Characteristics. Cell Rep 25, 1458–1468 e1454.

La Manno, G., Soldatov, R., Zeisel, A., Braun, E., Hochgerner, H., Petukhov, V., Lidschreiber, K., Kastriti, M.E., Lonnerberg, P., Furlan, A., et al. (2018). RNA velocity of single cells. Nature 560, 494–498.

Lavker, R.M., and Sun, T.T. (1982). Heterogeneity in epidermal basal keratinocytes: morphological and functional correlations. Science 215, 1239–1241.

Lavker, R.M., and Sun, T.T. (2000). Epidermal stem cells: properties, markers, and location. Proc Natl Acad Sci U S A 97, 13473–13475.

LeBoeuf, M., Terrell, A., Trivedi, S., Sinha, S., Epstein, J.A., Olson, E.N., Morrisey, E.E., and Millar, S.E. (2010). Hdac1 and Hdac2 act redundantly to control p63 and p53 functions in epidermal progenitor cells. Dev Cell 19, 807–818.

Lee, B., Villarreal-Ponce, A., Fallahi, M., Ovadia, J., Sun, P., Yu, Q.C., Ito, S., Sinha, S., Nie, Q., and Dai, X. (2014). Transcriptional mechanisms link epithelial plasticity to adhesion and differentiation of epidermal progenitor cells. Dev Cell 29, 47–58.

Li, J., Jiang, T.X., Hughes, M.W., Wu, P., Yu, J., Widelitz, R.B., Fan, G., and Chuong, C.-M. (2012). Progressive alopecia reveals decreasing stem cell activation probability during aging of mice with epidermal deletion of DNA methyltransferase 1. J. Invest. Dermatol. 132, 2681–2690.

Li, J., and Sen, G.L. (2015). Generation of Genetically Modified Organotypic Skin Cultures Using Devitalized Human Dermis. J Vis Exp, e53280.

Lim, X., and Nusse, R. (2013). Wnt signaling in skin development, homeostasis, and disease. Cold Spring Harb Perspect Biol 5.

Lim, X., Tan, S.H., Koh, W.L., Chau, R.M., Yan, K.S., Kuo, C.J., van Amerongen, R., Klein, A.M., and Nusse, R. (2013). Interfollicular epidermal stem cells self-renew via autocrine Wnt signaling. Science 342, 1226–1230.

Lin, C.H., Chiu, P.Y., Hsueh, Y.Y., Shieh, S.J., Wu, C.C., Wong, T.W., Chuong, C.-M., and Hughes, M.W. (2019). Regeneration of rete ridges in Lanyu pig (Sus scrofa): Insights for human skin wound healing. Exp. Dermatol. 28, 472–479.

Liu, Y., Lyle, S., Yang, Z., and Cotsarelis, G. (2003). Keratin 15 promoter targets putative epithelial stem cells in the hair follicle bulge. J Invest Dermatol 121, 963–968.

Lopez, R.G., Garcia-Silva, S., Moore, S.J., Bereshchenko, O., Martinez-Cruz, A.B., Ermakova, O., Kurz, E., Paramio, J.M., and Nerlov, C. (2009). C/EBPalpha and beta couple interfollicular keratinocyte proliferation arrest to commitment and terminal differentiation. Nat Cell Biol 11, 1181–1190.

Lopez-Pajares, V., Qu, K., Zhang, J., Webster, D.E., Barajas, B.C., Siprashvili, Z., Zarnegar, B.J., Boxer, L.D., Rios, E.J., Tao, S., et al. (2015). A LncRNA-MAF:MAFB transcription factor network regulates epidermal differentiation. Dev Cell 32, 693–706.

Lu, C.P., Polak, L., Rocha, A.S., Pasolli, H.A., Chen, S.C., Sharma, N., Blanpain, C., and Fuchs, E. (2012). Identification of stem cell populations in sweat glands and ducts reveals roles in homeostasis and wound repair. Cell 150, 136–150.

Lyle, S., Christofidou-Solomidou, M., Liu, Y., Elder, D.E., Albelda, S., and Cotsarelis, G. (1998). The C8/144B monoclonal antibody recognizes cytokeratin 15 and defines the location of human hair follicle stem cells. J Cell Sci 111 (*Pt 21*), 3179–3188.

Mackenzie, I.C. (1975). Ordered structure of the epidermis. J Invest Dermatol 65, 45–51.

Mackenzie, I.C. (1997). Retroviral transduction of murine epidermal stem cells demonstrates clonal units of epidermal structure. J Invest Dermatol 109, 377–383.

Macosko, E.Z., Basu, A., Satija, R., Nemesh, J., Shekhar, K., Goldman, M., Tirosh, I., Bialas, A.R., Kamitaki, N., Martersteck, E.M., et al. (2015). Highly Parallel Genome-wide Expression Profiling of Individual Cells Using Nanoliter Droplets. Cell 161, 1202–1214.

Mardaryev, A.N., Liu, B., Rapisarda, V., Poterlowicz, K., Malashchuk, I., Rudolf, J., Sharov, A.A., Jahoda, C.A., Fessing, M.Y., Benitah, S.A., et al. (2016). Cbx4 maintains the epithelial lineage identity and cell proliferation in the developing stratified epithelium. J Cell Biol 212, 77–89.

Mascre, G., Dekoninck, S., Drogat, B., Youssef, K.K., Brohee, S., Sotiropoulou, P.A., Simons, B.D., and Blanpain, C. (2012). Distinct contribution of stem and progenitor cells to epidermal maintenance. Nature 489, 257–262.

Mejetta, S., Morey, L., Pascual, G., Kuebler, B., Mysliwiec, M.R., Lee, Y., Shiekhattar, R., Di Croce, L., and Benitah, S.A. (2011). Jarid2 regulates mouse epidermal stem cell activation and differentiation. EMBO J 30, 3635–3646.

Mesa, K.R., Kawaguchi, K., Cockburn, K., Gonzalez, D., Boucher, J., Xin, T., Klein, A.M. and Greco, V. (2018). Homeostatic Epidermal Stem Cell Self-Renewal Is Driven by Local Differentiation. Cell Stem Cell 23, 677–686.

Mills, A.A., Zheng, B., Wang, X.J., Vogel, H., Roop, D.R., and Bradley, A. (1999). p63 is a p53 homologue required for limb and epidermal morphogenesis. Nature 398, 708–713.

Miyai, M., Hamada, M., Moriguchi, T., Hiruma, J., Kamitani-Kawamoto, A., Watanabe, H., Hara-Chikuma, M., Takahashi, K., Takahashi, S., and Kataoka, K. (2016). Transcription Factor MafB Coordinates Epidermal Keratinocyte Differentiation. J Invest Dermatol 136, 1848–1857.

Mok, K.W., Saxena, N., Heitman, N., Grisanti, L., Srivastava, D., Muraro, M.J., Jacob, T., Sennett, R., Wang, Z., Su, Y., et al. (2019). Dermal Condensate Niche Fate Specification Occurs Prior to Formation and Is Placode Progenitor Dependent. Dev Cell 48, 32–48 e35.

Moriyama, M., Durham, A.D., Moriyama, H., Hasegawa, K., Nishikawa, S., Radtke, F., and Osawa, M. (2008). Multiple roles of Notch signaling in the regulation of epidermal development. Dev Cell 14, 594–604.

Morris, R.J., Liu, Y., Marles, L., Yang, Z., Trempus, C., Li, S., Lin, J.S., Sawicki, J.A., and Cotsarelis, G. (2004). Capturing and profiling adult hair follicle stem cells. Nat Biotechnol 22, 411–417.

Mou, H., Vinarsky, V., Tata, P.R., Brazauskas, K., Choi, S.H., Crooke, A.K., Zhang, B., Solomon, G.M., Turner, B., Bihler, H., et al. (2016). Dual SMAD Signaling Inhibition Enables Long-Term Expansion of Diverse Epithelial Basal Cells. Cell Stem Cell 19, 217–231.

Nguyen, Q.H., Pervolarakis, N., Blake, K., Ma, D., Davis, R.T., James, N., Phung, A.T., Willey, E., Kumar, R., Jabart, E., et al. (2018). Profiling human breast epithelial cells using single cell RNA sequencing identifies cell diversity. Nat Commun 9, 2028.

Nijhof, J.G., Braun, K.M., Giangreco, A., van Pelt, C., Kawamoto, H., Boyd, R.L., Willemze, R., Mullenders, L.H., Watt, F.M., de Gruijl, F.R., et al. (2006). The cell-surface marker MTS24 identifies a novel population of follicular keratinocytes with characteristics of progenitor cells. Development 133, 3027–3037.

Nishio, H., Matsui, K., Tsuji, H., Tamura, A., and Suzuki, K. (2001). Immunolocalisation of the janus kinases (JAK)--signal transducers and activators of transcription (STAT) pathway in human epidermis. J Anat 198, 581–589.

Nusse, R., and Clevers, H. (2017). Wnt/beta-Catenin Signaling, Disease, and Emerging Therapeutic Modalities. Cell 169, 985–999.

Oh, J.W., Hsi, T.C., Guerrero-Juarez, C.F., Ramos, R., and Plikus, M.V. (2013). Organotypic skin culture. J Invest Dermatol 133, 1–4.

Ordovas-Montanes, J., Dwyer, D.F., Nyquist, S.K., Buchheit, K.M., Vukovic, M., Deb, C., Wadsworth, M.H., 2nd, Hughes, T.K., Kazer, S.W., Yoshimoto, E., et al. (2018). Allergic inflammatory memory in human respiratory epithelial progenitor cells. Nature 560, 649–654.

Oshimori, N., Oristian, D., and Fuchs, E. (2015). TGF-beta promotes heterogeneity and drug resistance in squamous cell carcinoma. Cell 160, 963–976.

Page, M.E., Lombard, P., Ng, F., Gottgens, B., and Jensen, K.B. (2013). The epidermis comprises autonomous compartments maintained by distinct stem cell populations. Cell Stem Cell 13, 471–482.

Philippeos, C., Telerman, S.B., Oules, B., Pisco, A.O., Shaw, T.J., Elgueta, R., Lombardi, G., Driskell, R.R., Soldin, M., Lynch, M.D., et al. (2018). Spatial and Single-Cell Transcriptional Profiling Identifies Functionally Distinct Human Dermal Fibroblast Subpopulations. J Invest Dermatol 138, 811–825.

Potten, C.S. (1974). The epidermal proliferative unit: the possible role of the central basal cell. Cell Tissue Kinet 7, 77–88.

Puram, S.V., Tirosh, I., Parikh, A.S., Patel, A.P., Yizhak, K., Gillespie, S., Rodman, C., Luo, C.L., Mroz, E.A., Emerick, K.S., et al. (2017). Single-Cell Transcriptomic Analysis of Primary and Metastatic Tumor Ecosystems in Head and Neck Cancer. Cell 171, 1611–1624 e1624.

Qiu, X., Mao, Q., Tang, Y., Wang, L., Chawla, R., Pliner, H.A., and Trapnell, C. (2017). Reversed graph embedding resolves complex single-cell trajectories. Nat Methods 14, 979–982.

Ramilowski, J.A., Goldberg, T., Harshbarger, J., Kloppmann, E., Lizio, M., Satagopam, V.P., Itoh, M., Kawaji, H., Carninci, P., Rost, B., et al. (2015). A draft network of ligand-receptor-mediated multicellular signalling in human. Nat Commun 6, 7866.

Rangarajan, A., Talora, C., Okuyama, R., Nicolas, M., Mammucari, C., Oh, H., Aster, J.C., Krishna, S., Metzger, D., Chambon, P., et al. (2001). Notch signaling is a direct determinant of keratinocyte growth arrest and entry into differentiation. EMBO J 20, 3427–3436.

Ren, J., Finney, R., Ni, K., Cam, M., and Muegge, K. (2019). The chromatin remodeling protein Lsh alters nucleosome occupancy at putative enhancers and modulates binding of lineage specific transcription factors. Epigenetics 14, 277–293.

Rompolas, P., Deschene, E.R., Zito, G., Gonzalez, D.G., Saotome, I., Haberman, A.M., and Greco, V. (2012). Live imaging of stem cell and progeny behaviour in physiological hair-follicle regeneration. Nature 487, 496–499.

Rompolas, P., Mesa, K.R., Kawaguchi, K., Park, S., Gonzalez, D., Brown, S., Boucher, J., Klein, A.M., and Greco, V. (2016). Spatiotemporal coordination of stem cell commitment during epidermal homeostasis. Science 352, 1471–1474.

Sada, A., Jacob, F., Leung, E., Wang, S., White, B.S., Shalloway, D., and Tumbar, T. (2016). Defining the cellular lineage hierarchy in the interfollicular epidermis of adult skin. Nat Cell Biol 18, 619–631.

Salzer, M.C., Lafzi, A., Berenguer-Llergo, A., Youssif, C., Castellanos, A., Solanas, G., Peixoto, F.O., Stephan-Otto Attolini, C., Prats, N., Aguilera, M., et al. (2018). Identity Noise and Adipogenic Traits Characterize Dermal Fibroblast Aging. Cell 175, 1575–1590 e1522.

Schiebinger, G., Shu, J., Tabaka, M., Cleary, B., Subramanian, V., Solomon, A., Gould, J., Liu, S., Lin, S., Berube, P., et al. (2019). Optimal-Transport Analysis of Single-Cell Gene Expression Identifies Developmental Trajectories in Reprogramming. Cell 176, 928–943 e922.

Segre, J.A., Bauer, C., and Fuchs, E. (1999). Klf4 is a transcription factor required for establishing the barrier function of the skin. Nat Genet 22, 356–360.

Sen, G.L., Reuter, J.A., Webster, D.E., Zhu, L., and Khavari, P.A. (2010). DNMT1 maintains progenitor function in self-renewing somatic tissue. Nature 463, 563–567.

Sen, G.L., Webster, D.E., Barragan, D.I., Chang, H.Y., and Khavari, P.A. (2008). Control of differentiation in a self-renewing mammalian tissue by the histone demethylase JMJD3. Genes Dev 22, 1865–1870.

Siebel, C., and Lendahl, U. (2017). Notch Signaling in Development, Tissue Homeostasis, and Disease. Physiol Rev 97, 1235–1294.

Smillie, C.S., Biton, M., Ordovas-Montanes, J., Sullivan, K.M., Burgin, G., Graham, D.B., Herbst, R.H., Rogel, N., Slyper, M., Waldman, J., et al. (2019). Intra- and Inter-cellular Rewiring of the Human Colon during Ulcerative Colitis. Cell 178, 714–730 e722.

Snippert, H.J., Haegebarth, A., Kasper, M., Jaks, V., van Es, J.H., Barker, N., van de Wetering, M., van den Born, M., Begthel, H., Vries, R.G., et al. (2010). Lgr6 marks stem cells in the hair follicle that generate all cell lineages of the skin. Science 327, 1385–1389.

Subramanian, A., Tamayo, P., Mootha, V.K., Mukherjee, S., Ebert, B.L., Gillette, M.A., Paulovich, A., Pomeroy, S.L., Golub, T.R., Lander, E.S., et al. (2005). Gene set enrichment analysis: a knowledge-based approach for interpreting genome-wide expression profiles. Proc Natl Acad Sci U S A 102, 15545–15550.

Tabib, T., Morse, C., Wang, T., Chen, W., and Lafyatis, R. (2018). SFRP2/DPP4 and FMO1/LSP1 Define Major Fibroblast Populations in Human Skin. J Invest Dermatol 138, 802–810.

Taniguchi, K., Arima, K., Masuoka, M., Ohta, S., Shiraishi, H., Ontsuka, K., Suzuki, S., Inamitsu, M., Yamamoto, K.I., Simmons, O., et al. (2014). Periostin controls keratinocyte proliferation and differentiation by interacting with the paracrine IL-1alpha/IL-6 loop. J Invest Dermatol 134, 1295–1304.

Teschendorff, A.E., and Enver, T. (2017). Single-cell entropy for accurate estimation of differentiation potency from a cell’s transcriptome. Nat Commun 8, 15599.

Ting, S.B., Caddy, J., Hislop, N., Wilanowski, T., Auden, A., Zhao, L.L., Ellis, S., Kaur, P., Uchida, Y., Holleran, W.M., et al. (2005). A homolog of Drosophila grainy head is essential for epidermal integrity in mice. Science 308, 411–413.

Tirosh, I., Izar, B., Prakadan, S.M., Wadsworth, M.H., 2nd, Treacy, D., Trombetta, J.J., Rotem, A., Rodman, C., Lian, C., Murphy, G., et al. (2016). Dissecting the multicellular ecosystem of metastatic melanoma by single-cell RNA-seq. Science 352, 189–196.

Trapnell, C., Cacchiarelli, D., Grimsby, J., Pokharel, P., Li, S., Morse, M., Lennon, N.J., Livak, K.J., Mikkelsen, T.S., and Rinn, J.L. (2014). The dynamics and regulators of cell fate decisions are revealed by pseudotemporal ordering of single cells. Nat Biotechnol 32, 381–386.

Trempus, C.S., Morris, R.J., Bortner, C.D., Cotsarelis, G., Faircloth, R.S., Reece, J.M., and Tennant, R.W. (2003). Enrichment for living murine keratinocytes from the hair follicle bulge with the cell surface marker CD34. J Invest Dermatol 120, 501–511.

Tumbar, T., Guasch, G., Greco, V., Blanpain, C., Lowry, W.E., Rendl, M., and Fuchs, E. (2004). Defining the epithelial stem cell niche in skin. Science 303, 359–363.

Tuncali, D., Bingul, F., Talim, B., Surucu, S., Sahin, F., and Aslan, G. (2005). Histologic characteristics of the human prepuce pertaining to its clinical behavior as a dual graft. Ann Plast Surg 54, 191–195.

Wang, S., Karikomi, M., MacLean, A.L., and Nie, Q. (2019). Cell lineage and communication network inference via optimization for single-cell transcriptomics. Nucleic Acids Res 47, e66.

Watt, F.M. (1983). Involucrin and other markers of keratinocyte terminal differentiation. J Invest Dermatol 81, 100s–103s.

Watt, F.M. (1998). Epidermal stem cells: markers, patterning and the control of stem cell fate. Philos Trans R Soc Lond B Biol Sci 353, 831–837.

Welsch, K., Holstein, J., Laurence, A., and Ghoreschi, K. (2017). Targeting JAK/STAT signalling in inflammatory skin diseases with small molecule inhibitors. Eur J Immunol 47, 1096–1107.

Wilson, S.I., Rydstrom, A., Trimborn, T., Willert, K., Nusse, R., Jessell, T.M., and Edlund, T. (2001). The status of Wnt signalling regulates neural and epidermal fates in the chick embryo. Nature 411, 325–330.

Wu, S.C. and Benavente, C.A. (2018). Chromatin remodeling protein HELLS is upregulated by inactivation of the RB-E2F pathway and is nonessential for osteosarcoma tumorigenesis. Oncotarget 66, 32580–32592.

Xu, M., Horrell, J., Snitow, M., Cui, J., Gochnauer, H., Syrett, C.M., Kallish, S., Seykora, J.T., Liu, F., Gaillard, D., et al. (2017). WNT10A mutation causes ectodermal dysplasia by impairing progenitor cell proliferation and KLF4-mediated differentiation. Nat Commun 8, 15397.

Yang, H., Adam, R.C., Ge, Y., Hua, Z.L., and Fuchs, E. (2017). Epithelial-Mesenchymal Micro-niches Govern Stem Cell Lineage Choices. Cell 169, 483–496 e413.

Zhang, Y., Tomann, P., Andl, T., Gallant, N.M., Huelsken, J., Jerchow, B., Birchmeier, W., Paus, R., Piccolo, S., Mikkola, M.L., et al. (2009). Reciprocal requirements for EDA/EDAR/NF-kappaB and Wnt/beta-catenin signaling pathways in hair follicle induction. Dev Cell 17, 49–61.

